# Single cell susceptibility to SARS-CoV-2 infection is driven by variable cell states

**DOI:** 10.1101/2023.07.06.547955

**Authors:** Sam Reffsin, Jesse Miller, Kasirajan Ayyanathan, Margaret C. Dunagin, Naveen Jain, David C. Schultz, Sara Cherry, Arjun Raj

**Affiliations:** Department of Bioengineering, School of Engineering and Applied Sciences, University of Pennsylvania, Philadelphia, PA, USA; Department of Pathology and Laboratory Medicine, University of Pennsylvania, Philadelphia, PA, USA; Genetics and Epigenetics, Cell and Molecular Biology Graduate Group, Perelman School of Medicine, University of Pennsylvania, Philadelphia, PA, USA; Department of Biochemistry and Biophysics, University of Pennsylvania, Philadelphia, PA, USA; Department of Microbiology, University of Pennsylvania, Philadelphia, PA, USA; Department of Genetics, Perelman School of Medicine, University of Pennsylvania, Philadelphia, PA, USA

## Abstract

The ability of a virus to infect a cell type is at least in part determined by the presence of host factors required for the viral life cycle. However, even within cell types that express known factors needed for infection, not every cell is equally susceptible, suggesting that our knowledge of the full spectrum of factors that promote infection is incomplete. Profiling the most susceptible subsets of cells within a population may reveal additional factors that promote infection. However, because viral infection dramatically alters the state of the cell, new approaches are needed to reveal the state of these cells prior to infection with virus. Here, we used single-cell clone tracing to retrospectively identify and characterize lung epithelial cells that are highly susceptible to infection with SARS-CoV-2. The transcriptional state of these highly susceptible cells includes markers of retinoic acid signaling and epithelial differentiation. Loss of candidate factors identified by our approach revealed that many of these factors play roles in viral entry. Moreover, a subset of these factors exert control over the infectable cell state itself, regulating the expression of key factors associated with viral infection and entry. Analysis of patient samples revealed the heterogeneous expression of these factors across both cells and patients *in vivo*. Further, the expression of these factors is upregulated in particular inflammatory pathologies. Altogether, our results show that the variable expression of intrinsic cell states is a major determinant of whether a cell can be infected by SARS-CoV-2.

## Introduction

Viruses hijack cellular machinery for entry and replication. The presence of this machinery at least in part determines the cellular tropism of the virus, generally defined as the cell type that is permissive to infection. These host factors include proteins and pathways required for each step in the viral life cycle including viral entry, viral processing, viral replication, and viral particle assembly and release (Chu et al., 2020; Puelles et al., 2020; Schneider-Schaulies, 2000; Tatsuo et al., 2000). However, even within a given cell type that is known to be infectable by a particular virus, only a subset of cells are infected (Heldt et al., 2015; Melms et al., 2021; Ravindra et al., 2021; Russell et al., 2018; Snijder et al., 2009; Snijder & Pelkmans, 2011).

We suggest that this specific subset of cells have higher levels of known host factors or express previously undescribed factors that promote susceptibility to viral infection. Hence, comparative profiling between infected and uninfected cells would in principle be a powerful method to identify factors that contribute to single cell differences in susceptibility within a permissive cell type. Such an approach represents an important complement to genetic screens, which can fail to detect factors for a number of reasons. Transcriptional profiling is typically done after infection, making it easier to identify which cells have been infected. However, since the infection itself modifies an infected cell’s transcriptional state, profiling cells after viral infection may not reveal the specific factors the cell expressed *prior* to infection.

Thus, while we know why some cell types are more permissive to infection than others, we have less knowledge of the within cell type differences that can affect susceptibility to infection. This is in part because it is relatively easy to identify the cell types that are infected by a virus independent of the detection of the viral infection itself, as compared to the identification of the individual infectable cells within a given cell type. For instance, ACE2, the receptor that SARS-CoV-2 uses to enter the cell, is expressed in respiratory epithelial club and ciliated cells, and these cell types are indeed the most susceptible to infection (Melms et al., 2021; Ravindra et al., 2021). However, during a bona fide infection, only a subset of club and ciliated cells are infected, suggesting that the expression of ACE2 alone may not be sufficient to explain why some cells are susceptible to the virus (Ravindra et al., 2021).

Cell culture models are used to determine the host pathways and players required for infection. However, even in these highly controlled settings, seemingly homogeneous cells can show differences in viral susceptibility. For instance, the Calu-3 cell line (an adenocarcinoma lung cell line) is a well-established model for infection of the human respiratory epithelium with SARS-CoV-2, as these cells endogenously express ACE2 and TMPRSS2, two essential factors for viral entry. Under limiting conditions, infection of Calu-3 cells with SARS-CoV-2 is variable, recapitulating the conditions that cells are under after initial exposure to virus *in vivo.* While it is possible that extrinsic factors could be responsible for this variability (Russell et al. 2018; Belser et al. 2022), it is also possible that the subset of cells that are infectable under such conditions are in a distinct cell state that is highly susceptible to infection. Identifying the specific factors that contribute to this single cell susceptibility is particularly challenging for SARS-CoV-2 infection because SARS-CoV-2 rapidly shuts off host cell transcription both *in vitro* and *in vivo* (Acheampong et al., 2022; Burke et al., 2021; Finkel et al., 2021; Thoms et al., 2020). Therefore, determining the transcriptional differences that correspond to variable susceptibility to SARS-CoV-2 infection is not possible using post-facto profiling approaches.

We thus set out to profile these highly susceptible cells using methods that can report on the state of cells that are destined for infection *prior to infection with the virus*. A number of such retrospective methods relying on DNA barcoding have been developed recently (Biddy et al., 2018; Emert et al., 2021; Goyal et al., 2021; Jain et al., 2023; Weinreb et al., 2020), but they have not yet been applied to the question of heterogeneity in viral infection. Such methods have the potential to reveal novel cell subpopulations and associated factors that may be critical for initial susceptibility to infection.

Here, we used a retrospective single cell method (Emert et al., 2021) to demonstrate that particular cells within a population are intrinsically more likely to become infected with SARS-CoV-2. The identified population of highly susceptible cells are a subset of ACE2 expressing cells that were further enriched for high expression of *TIG1*. Genetic knockout of factors expressed within the *TIG1*-high state led to decreased viral infection, demonstrating functional roles for these factors in the SARS-CoV-2 lifecycle. We identified known factors that promote infection such as *AXL* (Wang et al., 2021) and *TSPAN8* (Hysenaj et al., 2021). In addition, we identified previously unknown factors such as *TIG1* that altered the cell’s intrinsic susceptibility to infection by manipulating the expression of factors implicated in viral entry. Analysis of patient data suggested the existence of these specific and heterogeneously expressed cell states in the human lung, potentially pointing to a role for the identified factors in determining which epithelial cells are most permissive to infection *in vivo*.

## Results

### Single cell clone tracing identifies host cell gene expression states that are highly susceptible to infection with SARS-CoV-2

We set out to determine whether variability in host gene expression within single cells could affect the likelihood of a cell becoming infected. To do so, we used the human respiratory Calu-3 cell line, which can be infected by SARS-CoV-2 and recapitulates many key features of lung epithelial biology making it a useful model for infection. At a low multiplicity of infection, we found that 1.5% of the cells 24 hours post infection and 12.5% of the cells 48 hour post infection were infected as measured by single molecule RNA FISH against viral genomic RNA (Figure 1A, Figure S4D-E). We initially determined whether we could simply profile infected and uninfected cells post-infection (Fig. S1A). However, as previously observed, we found that the virus rapidly altered transcription in the host cell, leading to the loss of expression of housekeeping genes specifically in infected cells, rendering such comparisons difficult to interpret (Figure S1B). This further motivated the need for retrospective identification and profiling of cells that are destined to be infected before infection occurs.

**Figure 1:**
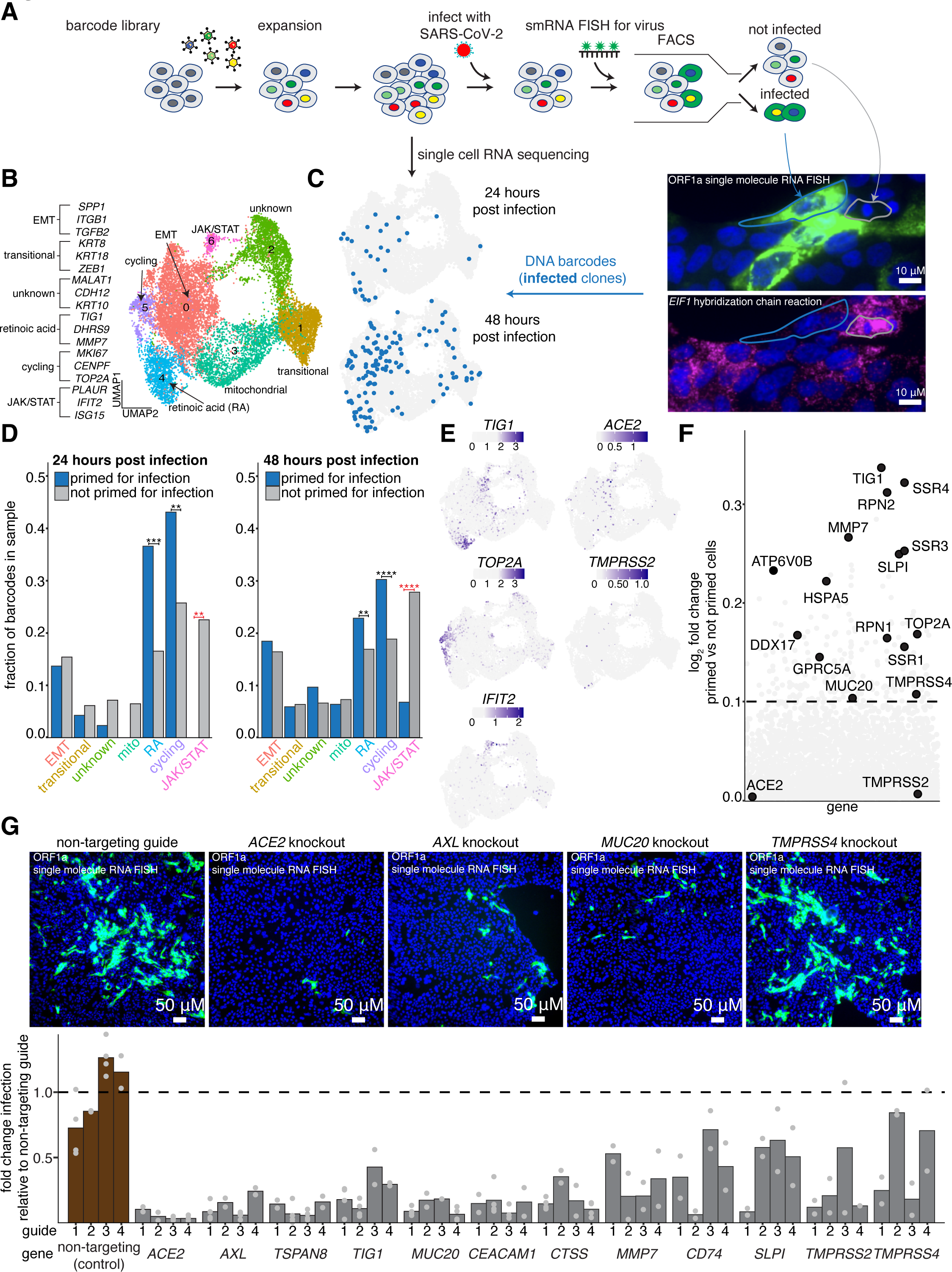
Single cell clone tracing reveals a distinct host transcriptional state that is highly susceptible to infection with SARS-CoV-2. **A.** Schematic of Rewind approach for retrospectively identifying transcriptional states of single Calu-3 cells that are highly susceptible to infection. We retrovirally transduced 100,000 Calu-3 cells at an MOI of ∼0.1. After 7 days (∼3 population doublings), we split clones into two arms, performing 10x single cell RNA sequencing on one arm and infecting the other arm with SARS-CoV-2. At both 24 hours and 48 hours post infection, the infected arm was fixed and FACS sorted on viral single molecule RNA FISH signal (FISH probes targeting ORF1a within the viral genome, see methods). Clones with infected and not infected barcodes were then annotated in the matched pre-infection snapshot single cell RNA sequencing data. **B.** UMAP plot showing transcriptional clusters in starting Calu-3 population. Cluster labels were assigned based on expression of marker genes within each cluster. **C.** Location in UMAP space of barcodes of cells primed for infection collected at each time point (n=29 clones at 24 hours post infection, n = 132 clones at 48 hours post infection). **D.** The fraction of primed barcodes found in each cluster for each time point collected. To correct for the different number of cells within each cluster, the fractions were normalized to the number of total recovered barcodes within each cluster. Significance was determined by comparing the actual barcode enrichment of primed and not primed cells within each cluster to a randomly sampled distribution of the total recovered barcodes (n=100 simulations, see methods for details). One star represents a p-value < 0.05. Two stars represent a p-value < 0.01. Three stars represent a p-value < 0.001. Four stars represent a p-value < 0.0001. **E.** Normalized expression of select marker genes from each cluster that is enriched for primed barcodes or enriched for not primed barcodes. **F.** Direct comparison of gene expression profiles of cells primed for infection compared to those that were not. **G.** To validate that our identified markers promote SARS-CoV-2 infection, we used CRISPR-Cas9 to knockout a subset of genes that were found to be enriched in cells primed for infection. Briefly, Calu-3 were transduced with a guide construct containing both a sgRNA against a target gene and the Cas9 transgene. For each gene, we tested four independent guides. After transduction, cells were put under puromycin selection for 8 days, then infected with SARS-CoV-2. 48 hours post infection, cells were fixed and analyzed for viral single molecule RNA FISH signal via microscopy. All data is normalized to non-targeting control guides against the AAVS1 locus. Bars shown are the average of two independent biological replicates, except for certain guides targeting the non-targeting control, *ACE2, AXL, CEACAM1, MUC20, TIG1,* and *TSPAN8* for which the bars represent the average of four independent biological replicates.

To retrospectively identify the individual cells that were most likely to become infected, we applied a modified version of our previously described Rewind methodology (Emert et al., 2021). Briefly, Rewind uses lentiviral-based DNA barcodes that are integrated into the cell’s genome and expressed as mRNA in the 3’ UTR of a GFP transgene. Since integrated lentiviruses are heritable through cell divisions, these barcodes can be used to connect single cells across time. Once cells are transduced with the barcode, “twins”, or cells with a recent common progenitor, can be collected as the cells divide, serving as snapshots of the current state of the cell. If twins are separated into two groups—one that is immediately profiled to obtain a baseline transcriptional state, and another that is infected, sorted on viral RNA signal, and subsequently sequenced for barcode abundance—the overlapping barcodes found across the different groups can be used to connect infection to the original transcriptional state of the cell (Figure 1A).

An important assumption underlying the use of Rewind is that a cell’s propensity to become infected is at least partially intrinsic to the cell (i.e., not fully dependent on external factors, such as local environment) and heritable through at least a few cell divisions. To test this assumption, we first barcoded Calu-3 cells using our Rewind approach, targeting a low multiplicity of infection to ensure that the majority of cells had just one integration event. After allowing the cells to undergo a set number of divisions (creating multiple “clones” per barcode lineage), the clones were split into two experimental arms and independently infected with SARS-CoV-2 (Figure S2A). If there were an intrinsic component to infection, we would expect a high degree of overlap in the barcodes that are recovered from infected cells across each experimental arm (Goyal et al., 2021; Jain et al., 2023). Conversely, if infection were largely driven by other differences independent of heritable factors intrinsic to the cell, the number of overlapping barcodes would be much lower, as predicted by chance. We found a high degree of overlap in the barcodes isolated from infected cells between split cultures. In the first replicate, we found 55.2% of the barcodes in split 1 and 35.9% of the barcodes in split 2 overlapping, compared to an expected random overlap of 0.14% and 0.09%, respectively. In the second replicate, we found 59.8% of the barcodes in split 1 and 62.7% of the barcodes in split 2 overlapping, compared to an expected random overlap of 0.31% and 0.33%, respectively. (Figure S2B-C). Thus, infectability was driven to a large extent by cell-intrinsic differences that persist over at least a few cell divisions.

Given that we observed significant intrinsic memory for a cell to be susceptible to infection, we then applied Rewind to retrospectively determine what transcriptional features were associated with these highly susceptible cells. We barcoded Calu-3 cells and after three cell divisions, twins for each clone were split into two arms. Single cell RNA sequencing (10x Chromium platform V3) was performed on one set to obtain a snapshot of cellular states before infection. The second set was infected with SARS-CoV-2. From this arm, we used single molecule RNA FISH (smFISH) against the SARS-CoV-2 genome (ORF1a, see methods) coupled with FACS to purify infected and uninfected cells at either 24 or 48 hours post infection. We next extracted the genomic DNA, amplified, and then sequenced the barcodes from each collected population. We further matched barcodes present in either infected or uninfected bystander cells with barcodes identified in the single cell RNA sequencing data from the pre-infection snapshot (Figure 1A, Figure S3A).

### Calu-3 cells comprise diverse cellular states

We first explored the baseline transcriptional heterogeneity in Calu-3. After pre-processing and clustering of our data (Seurat v3), we broadly identified two populations: those that expressed mucins such as *MUC5AC* and *MUC5B* and those that expressed more traditional epithelial cell markers such as *EPCAM* and various keratins (Figure S4B). Within *MUC5AC-*high cells, we found a subtype featuring retinoic acid associated genes (*TIG1, DHRS9, BMP7, LTBP2)* (Nagpal et al., 1996; C. Wang et al., 2011) as well as subtypes that featured genes associated with cell cycle (*TOP2A, MKI67, CENPF)*, epithelial-mesenchymal transition (EMT) (*SPP1, TGFB2, ITGB1, VCAN)*, and JAK/STAT signaling (*PLAUR, ISG15, IFIT2)*. Within the more traditional epithelial cells, one subpopulation expressed genes known to be involved with type-2 pneumocyte differentiation in the lung epithelium (*KRT8, KRT18, ZEB1)* (Jiang et al., 2020; Strunz et al., 2020; Verheyden & Sun, 2020) (Figure 1B, Figure S4C). Given the importance of *ACE2* and *TMPRSS2* in SARS-CoV-2 infection, we wanted to determine if any cluster was enriched for their expression. We found very few cells with detectable expression of either factor, potentially due to low levels of expression of these genes at the single cell mRNA level (Figure 1E). Therefore, at baseline, individual Calu-3 cells have distinct transcriptional states.

### Expression differences observed in cells primed for infection

We then classified cells from the pre-infection arm as primed or not primed based on whether the cell’s barcode was also found in either the infected or uninfected group respectively after infection (Figure 1C, see methods). After performing quality control on the recovered barcodes, we identified 29 infected and 1096 uninfected bystander clones at 24 hours post infection. The infection rate calculated from these barcode numbers is 2.6%, which is comparable to the infection rate of 1.5% as determined by single molecule RNA FISH. At 48 hours post infection, we identified 132 infected and 598 uninfected clones whose twins were infected and uninfected (resulting in an infection rate of 18.1%), again comparable to the actual infection rate of 12.5% obtained by single molecule RNA FISH (Figure S4D-E). The increased rate of infection at this time point was consistent with bystander effects and secondary infections, where cells that were not initially infected may activate new transcriptional responses due to changes that occur as a result of nearby infected cells (Bost et al., 2020; Hancock et al., 2018; Steuerman et al., 2018; Zanini et al., 2018).

Next, we mapped the barcodes of the identified infected and uninfected bystander cells to the transcriptome clusters we obtained from the baseline Calu-3 population. We calculated the proportion of barcodes present in each cluster, taking into account barcode recovery statistics (see methods, Figure S3B-C). At both 24 and 48 hours post infection, we found that two clusters were enriched for cells that were primed for infection: one that corresponded to fast-cycling cells marked by the expression of *TOP2A, CENPF,* and *MKI67*, and the other marked by retinoic acid signaling, in particular *TIG1* (Figure 1D-E). Beyond *TIG1*, the retinoic acid responsive cluster also expressed genes related to extracellular matrix remodeling (*MMP7)* (Huntington et al., 2004; Zhang et al., 2017), NF-kappaB regulation (*CEACAM1, SLPI)* (Ellerbeck et al., 2011; Gencheva et al., 2010), and Met signaling *(MUC20)* (Higuchi et al., 2004).

We also identified clusters that had a significant depletion of primed cells. Interestingly, we saw a lack of primed cells in the JAK/STAT signaling cluster (Figure 1D). This cluster expressed various dsRNA sensors (*RIGI*, *OASL*, *OAS2*) and interferon stimulated genes (*IFIT2*, *ISG15*, *MX1*), suggesting that there is a subset of cells that have high expression of this signaling pathway compared to the population average prior to infection, likely conferring intrinsic protection from viral infection.

While primed cells were enriched in particular clusters, those clusters nevertheless contained cells that were not primed, and some primed cells were present in other clusters. We next assessed whether there were additional genes that were over or under-represented within primed cells that were not captured in our cluster analysis. We performed an alternative “pseudobulk” analysis in which we averaged together the transcriptomes of all primed cells in one group and non-primed cells in another and performed standard differential expression analysis. Comparing primed cells identified 24 hours post infection to unprimed cells, the set of genes that were differentially expressed was virtually identical to those identified by our cluster analysis (Figure 1F, Figure S4F). We next compared the primed and unprimed cells identified 48 hours post infection. We found fewer differentially expressed genes, although *TIG1* and *MMP7* still emerged as top hits (Figure S4F). Beyond *TIG1* and *MMP7*, the genes that we did find were noticeably less enriched for a particular cluster, and were largely housekeeping factors including ribosomal genes. These findings supported the notion that genes associated with priming early in infection were more likely to be drivers of susceptibility, and so we focused on these factors for further analysis.

### Validation of factors that mark cells highly susceptible to infection

To validate the association between the expression of the identified genes and permissivity to infection, we adapted the Rewind approach to include a sorting step to pre-enrich for particular populations of cells. As before, after barcoding cells and allowing for three cell divisions, we split the cells into two groups. We infected one group with SARS-CoV-2. In the second group, we sorted fixed cells that were labeled for *TIG1* and *MMP7* expression using Hybridization Chain Reaction v3 (an amplified RNA FISH technique that increases signal intensity to accurately sort on a flow cytometer) (Choi et al., 2018). We isolated four populations based on high or low expression of these two genes. After sorting the populations, we extracted genomic DNA and recovered barcodes from high and low *TIG1* and *MMP7* expressing cells (Figure S5A). As a control for potential artifacts associated with sorting, we also sorted cells based on high and low expression of the housekeeping gene *EIF1*, with the number of cells in each *EIF1* bin carefully matched to their high and low *TIG1* and *MMP7* counterparts to account for potential technical artifacts related to the rate of barcode recovery. We then calculated the percentage of barcodes in each sorted population that overlapped with the barcodes recovered from the infected cells; a high degree of overlap would indicate that cells with high (or low) expression of a given gene have a greater propensity for infection compared to the bulk population (Figure S5A). We found that cells with high *TIG1* and high *MMP7* expression had an increased proportion of overlapping barcodes with the infected cells than the housekeeping control (83.3% for *TIG1*-high cells and 37.5% for *MMP7*-high cells, compared to 22.75% *EIF1*-high cells in replicate 1; 45.2% and 25% respectively, compared to 10.4% in replicate 2) (Figure S5B), thus validating that the Rewind experiments faithfully revealed a gene expression program associated with susceptibility to infection.

*TIG1* was originally identified as a gene induced by the retinoid tazarotene (Nagpal et al., 1996). Therefore, we pre-treated cells with tazarotene to induce *TIG1* and then infected the cells with SARS-CoV-2. Our results showed that the treatment of Calu-3 cells with tazarotene led to a variable induction of *TIG1* across diverse conditions. Across all conditions, we found that the percentage of cells expressing *TIG1* prior to infection strongly correlated with the percentage of cells infected with SARS-CoV-2 at 48 hours post-infection (correlation coefficient of R=0.91, p-value = 4.8e-13) (Figure S5C). Altogether, when *TIG1* expression increased, so too did infection, thus supporting the finding that *TIG1* serves as a marker for a permissive cell state to SARS-CoV-2 infection across a range of conditions.

### Intrinsic factors drive the spatial organization of Calu-3 cells

Calu-3 cells grow as islands, and we observed that the majority of cells infected with SARS-CoV-2 were present at the boundaries of cellular islands, a finding reported by other groups (Snijder et al., 2009). We also observed high expression of primed state markers (e.g. *MMP7*, *TIG1*) at these boundaries (Figure S4G). Given that a cell and its twin are plated independently, how can a cell and its twin both have a higher likelihood of infection when there is no guarantee that the twins would end up at a boundary in both arms of the experiment? In other words, how can the primed state for infection be an intrinsic property of the cell, but also be affected by seemingly extrinsic properties, such as location at a spatial boundary? To address this paradox, we performed live cell imaging, tracking the process by which Calu-3 cells attach and organize after seeding. We found that the boundaries of aggregates were not extrinsically defined based on where cells attached to the dish, but rather that the boundaries themselves were formed by the behavior of particular cells (Video S1-2). Cells fated for boundaries formed small holes in the monolayer, which they then expanded. As such, these cells were located at the boundary precisely because they manipulate the other cells in the culture to form these boundaries. Altogether, these boundary cells appear to be intrinsically programmed.

### Cells that are primed for infection likely represent a subset of ACE2 expressing cells

Viral tropism is at least in part defined by the expression of the cellular receptor essential for infection. Because the receptor ACE2 is required for SARS-CoV-2 infection, we set out to determine if the variability in infection across cells reflects variability in the expression of ACE2. Thus, cells identified as primed for infection may simply represent the highest ACE2 expressing cells in the population. Therefore, we characterized the single cell variability of ACE2 across Calu-3 cells. We first mined our Rewind dataset to determine the relative expression of *ACE2* in primed and not primed cells. However, as mentioned previously, we found very few cells with detectable *ACE2* expression in these data across all clusters (n=17 cells out of 14,057 with more than one UMI count assigned to *ACE2*), at least in part due to the low level expression of *ACE2* mRNA. Single molecule RNA FISH provided a more sensitive quantification, showing that 1.97% of cells had at least 10 detected transcripts (cutoff determined by the histogram in Figure S6H), with a mean *ACE2* transcript count per cell of 1.15 (Figure S6G).

Given how low and variable *ACE2* mRNA transcript abundance was and the fact that ultimately SARS-CoV-2 relies on ACE2 protein for infection, we used immunofluorescence to measure ACE2 protein levels across individual cells. We confirmed the specificity of our antibody by observing a ∼4-fold reduction in signal intensity when *ACE2* was knocked out (Figure 3F). We found that a much greater percentage (approximately 9%) of cells had high levels of ACE2 protein (Figure 2A-B). This percentage was also much higher than the number of cells that were initially infected (1.5% 24 hours post infection, Figure S4D). The finding that the percentage of initially infected cells was lower than the percentage of high-expressing ACE2 cells suggests that only a subset of the ACE2 expressing cells in the population were primed for infection.

**Figure 2:**
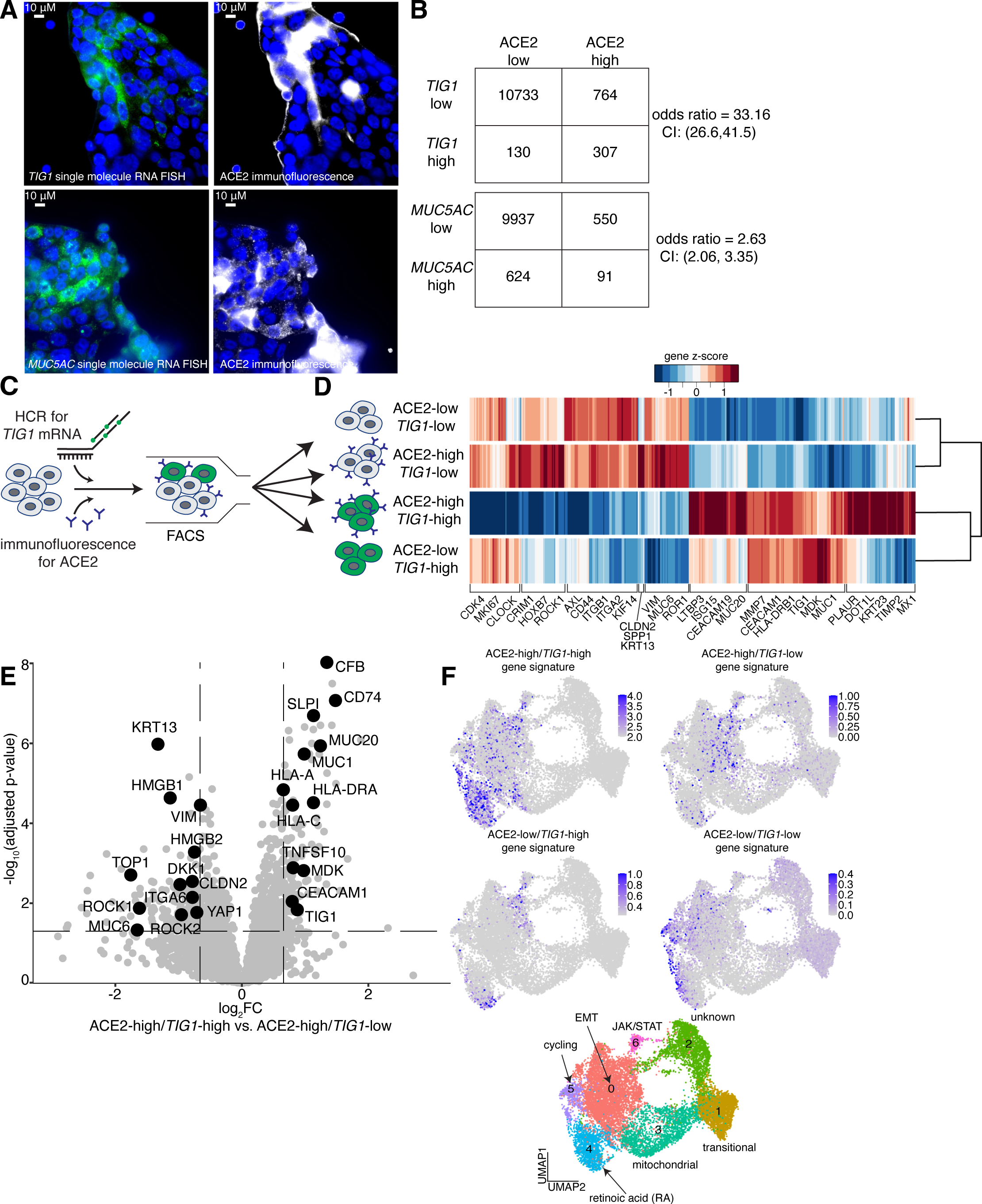
Calu-3 identified as primed for infection represent a subset of ACE2 expressing cells. **A.** Immunofluorescence against ACE2 protein and single molecule RNA FISH against *TIG1* mRNA or *MUC5AC* mRNA was performed simultaneously in the same cells. **B.** Contingency tables representing the various cell count frequencies of the identified populations from **A.** Odds ratios were calculated and confidence intervals were assigned using the Fisher’s Exact Test. **C.** Experimental design to isolate various ACE2 and *TIG1* expressing populations for differential expression analysis. **D.** Heatmap of differentially expressed genes (logFC > 1, p-value < 0.05) from each population. For each gene, a z-score was calculated relative to the average expression of that gene across all samples. If a gene was expressed below the average in a given sample, that gene’s z-score is reported as a negative value. **E.** Volcano plot comparing gene expression of ACE2-high/*TIG1*-high Calu-3 to ACE2-high/*TIG1-*low Calu-3. A representative subset of differentially expressed genes were selected and highlighted in black. **F.** The gene signatures identified in **C.** were mapped onto the Rewind single cell RNA sequencing dataset using the AddModuleScore function in Seurat v3. The score of each gene signature in each cell is plotted in UMAP space.

**Figure 3:**
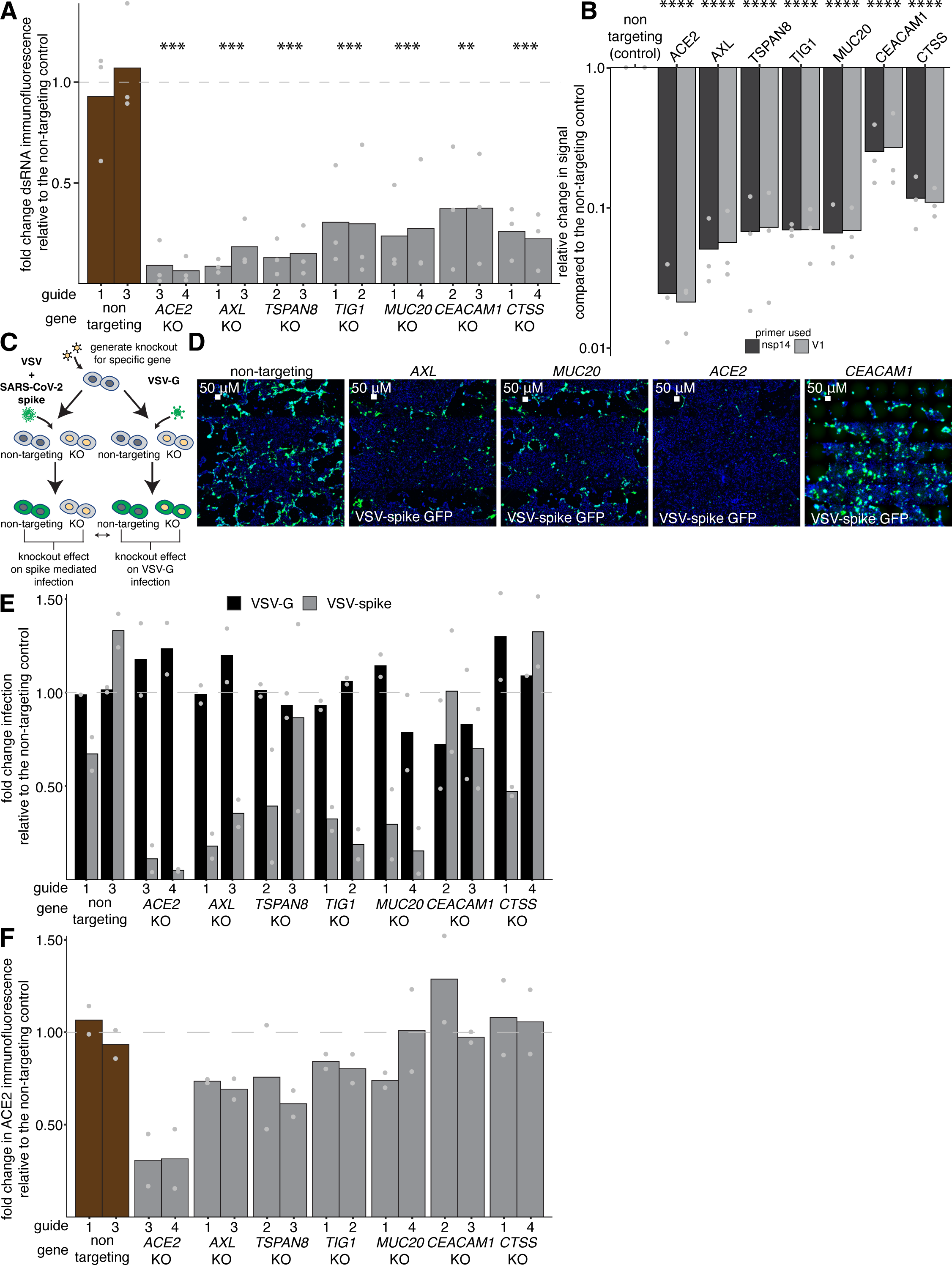
Factors expressed in the TIG1-high state regulate infection by diverse modes of action. **A.** Quantification of the percentage of cells infected as determined by immunofluorescence signal against viral dsRNA (J2) across each knockout condition compared to the non-targeting control. Each bar is the mean of three independent biological replicates. The top two performing guides for each gene from previous experiments were used. Significance was determined by performing a t-test on the average fold change for each gene relative to the non-targeting control. Two stars represent a p-value < 0.01. Three stars represent a p-value < 0.001. **B.** Relative change in viral RNA detected by qPCR for two different primer sets. For each gene, the same two guides as **A** were pooled together for subsequent analysis. Each bar is the average of three independent biological replicates. Significance was determined by performing a t-test on the average fold change in signal for each gene across primers relative to the non-targeting control. Four stars represent a p-value < 0.0001. **C.** Schematic for SARS-CoV-2 spike entry assay. Briefly, knockout cells were generated using CRISPR-Cas9. Cells for each knockout were then infected with either VSV-spike, featuring the SARS-CoV-2 spike protein in place of the native VSV glycoprotein, or VSV-G containing its wild type glycoprotein. 24 hours post infection, cells were fixed and analyzed via high throughput imaging for a change in the percent of cells infected by each virus. **D.** Representative images showing VSV-spike fluorescent signal across multiple knockout conditions. From left to right; non-targeting control, *AXL* knockout, *MUC20* knockout, *ACE2* knockout, and *CEACAM1* knockout. Viral signal is labeled by GFP (green). Cell nuclei are labeled by DAPI (blue). **E.** Quantification of the percent of cells infected by VSV-G signal (black) and VSV-spike signal (gray) across knockout conditions. Signal is normalized to the non-targeting control for each virus. Each bar is the average across two biological replicates. **F.** Quantification of ACE2 immunofluorescence signal across various knockout conditions. Signal is normalized to the non-targeting control. Each bar is the average across two biological replicates.

Given that our Rewind experiments identified *TIG1* as a marker of priming, we next measured expression of both *TIG1* (smRNA FISH) and ACE2 (immunofluorescence) simultaneously in Calu-3 cells by microscopy. Our results showed that while ACE2 protein and *TIG1* expression were strongly correlated (odds ratio of 33.16, confidence interval 26.6, 41.5), only a subset of ACE2-high cells were also *TIG1*-high (Figure 2B). Specifically, 28.7% of the ACE2-high cells detected also expressed high levels of *TIG1*, while 70.3% of *TIG1*-high cells were high for ACE2. To ensure that the correlation between *TIG1* and ACE2 was not an artifact of fluorescent bleedthrough due to high antibody signal or autofluorescence, we also measured *MUC5AC* mRNA and ACE2 protein using the same fluorescent dyes. The correlation between *MUC5AC* and ACE2 was significantly weaker (odds ratio of 2.63, confidence interval 2.06, 3.35), with only 14.2% of ACE2-high cells being high for *MUC5AC* and 12.7% of *MUC5AC*-high cells staining high for ACE2 (Figure 2B). These results suggest that additional factors beyond ACE2 expression levels determine which cells are most susceptible to infection.

We then queried how these ACE2 and *TIG1* subpopulations mapped to the Calu-3 clusters we previously observed. We stained for ACE2 protein and *TIG1* mRNA using RNA FISH (HCRv3) and performed FACS purification to sort cells that were high for both ACE2 protein and *TIG1* mRNA, low for both ACE2 protein and *TIG1* mRNA, or high for one or the other only, and then sequenced each population’s mRNA in bulk to identify differentially expressed genes (Figure 2C). Using the gene signatures that mark each of the sorted populations, we then mapped these signatures to our single cell RNA sequencing data to identify which subset of Calu-3 cells were the closest match to these populations (Seurat v3 AddModuleScore function, see methods). Our analysis revealed that the gene signature from the ACE2-high and *TIG1*-high population overlapped with cells in the *TIG1-*high state which are primed for infection (Figure 2D-F). Additional genes in this cluster include *MUC20*, *CEACAM1*, and *MMP7.* In contrast, cells that were high for ACE2 protein and low for *TIG1* mRNA expression overlapped with the EMT cluster and included an EMT signature (*SPP1, AXL*, *ITGA2*, *ITGB1*, *VIM*), and stem-like epithelial cell markers (*KRT13*, *ROCK1*, *ROCK2*, *MUC6*) (Hewitt & Lloyd, 2021; Montoro et al., 2018; Yin et al., 2022; Zhou et al., 2022) (Figure 2D-F). Irrespective of *TIG1* expression, cells that had low levels of ACE2 protein appeared to be cycling and were marked by the expression of *MKI67, TOP2A*, and *CDK4.* Overall, these data suggest that cells in a primed state for infection represent a subset of ACE2-high cells and that ACE2 expression alone is not sufficient to identify the population of cells that are highly susceptible to infection.

### Genes associated with priming promote susceptibility to infection

The identification of a specific cellular state associated with infection raised the possibility that the factors expressed within this state included host factors that specifically promote infection. To assess the functional role of these factors, we used CRISPR-Cas9 to knock out a subset of genes that were highly expressed in the *TIG1*-high population, as well as *ACE2* and *TMPRSS2* as positive controls (Figure 1G, Figure S6A-G). We used smRNA FISH against the viral genome to quantify infection. Compared to the non-targeting control (AAVS1), knockout of *AXL*, *CEACAM1*, *CTSS*, *MUC20*, *TIG1*, and *TSPAN8* all reduced infection while *CD74*, *SLPI*, and *TMPRSS4* did not (Figure 1G). Both *AXL* and *TSPAN8* have been previously implicated as factors involved with SARS-CoV-2 infection (Bohan et al., 2021; Hysenaj et al., 2021; Wang et al., 2021), while *CEACAM1* is a receptor for mouse hepatitis virus that has not been shown to have a role in SARS-CoV-2 infection (Williams et al., 1991). In addition, *MUC20* expression has been associated with severe COVID-19 disease *in vivo*, where its expression was shown to be depleted in patients with severe disease relative to those with mild disease (Smet et al., 2021). We further validated the role these genes played in infection using qPCR to quantify viral RNA and immunofluorescence against viral dsRNA (Figure 3A-B). Thus, Rewind identified a cellular state that promotes infection as well as host factors whose expression directly affects susceptibility to infection with SARS-CoV-2.

### Priming-associated host factors promote viral entry

Next, we set out to determine how these factors impact SARS-CoV-2 infection. We first tested whether these factors were involved in viral entry using a replication-competent vesicular stomatitis virus (VSV) that either expresses its native entry glycoprotein G or the SARS-CoV-2 spike glycoprotein (Case et al., 2020). This recombinant VSV also expresses GFP, allowing us to directly measure infection by microscopy. If a host factor is required for entry via SARS-CoV-2 spike glycoprotein, removal of that factor would reduce the infection of VSV-Spike but not VSV-G. Thus, by comparing the requirement of a host factor for infection by SARS-CoV-2, VSV-G, and VSV-Spike, we can determine if a host factor specifically impacts Spike-mediated entry (Figure 3C). We deleted each factor and challenged the cells with VSV-G or VSV-Spike. We found that none of the genes tested had an effect on VSV-G infection, and that knockout of the known entry receptor ACE2 showed a decrease in VSV-spike infection but not VSV-G. Furthermore, we found that knocking out *AXL*, *MUC20*, and *TIG1* all reduced viral infection by VSV-Spike, while *CEACAM1* and *CTSS* knockout had no effect (Figure 3D-E). These data suggest that *AXL*, *MUC20* and *TIG1* promote SARS-CoV-2 entry.

We then tested if the decrease in viral entry caused by knocking out *AXL*, *MUC20*, or *TIG1* was potentially due to changes in ACE2 protein levels. We performed immunofluorescence to measure ACE2 in each knockout condition. Our results revealed minor changes in ACE2 protein levels across the knockouts, indicating that the impact of *AXL*, *MUC20*, and *TIG1* on viral entry was not mediated by the regulation of ACE2 protein (Figure 3F). Of note, AXL has previously been shown to facilitate viral entry through the binding of viral glycoproteins (Wang et al., 2021).

### Genetic perturbations reveal modulators of a cell state that is enriched for viral entry

The host factors we identified included several molecules that could in principle affect viral infection either by direct interaction with the virus during some part of its life cycle or by influencing the expression of such factors (or both). As a starting point, we used pseudotime analysis to generate a putative temporal ordering of the expression of genes as Calu-3 cells transition through the various states identified by single cell RNA sequencing. In order to calculate trajectories, we performed additional single cell RNA sequencing on Calu-3 cells to obtain increased sequencing depth. We observed similar clusters as our original Rewind data, including subsets of cells that were in cycling, EMT, JAK/STAT, and retinoic acid responsive clusters (Figure S7A-B). We then fit pseudotime trajectories to our single cell RNA sequencing data to determine if the factors identified by Rewind that we found to promote infection drive these orderings (Slingshot, see methods).

We found three distinct lineages starting from a subset of cycling cells with stem-like regulatory network signatures marked by *HES1*, *POU2F1*, and *NFYB* (Moriyama et al., 2006; Ohtsuka et al., 2001; Shen et al., 2017) (Figure 4A-B, Figure S7C). The first trajectory appears to be cells that continue to cycle, whereas the second trajectory ends at the *TIG1*-high cluster which represents cells primed for infection, and the third ends at the JAK/STAT cluster (Figure 4B).

**Figure 4:**
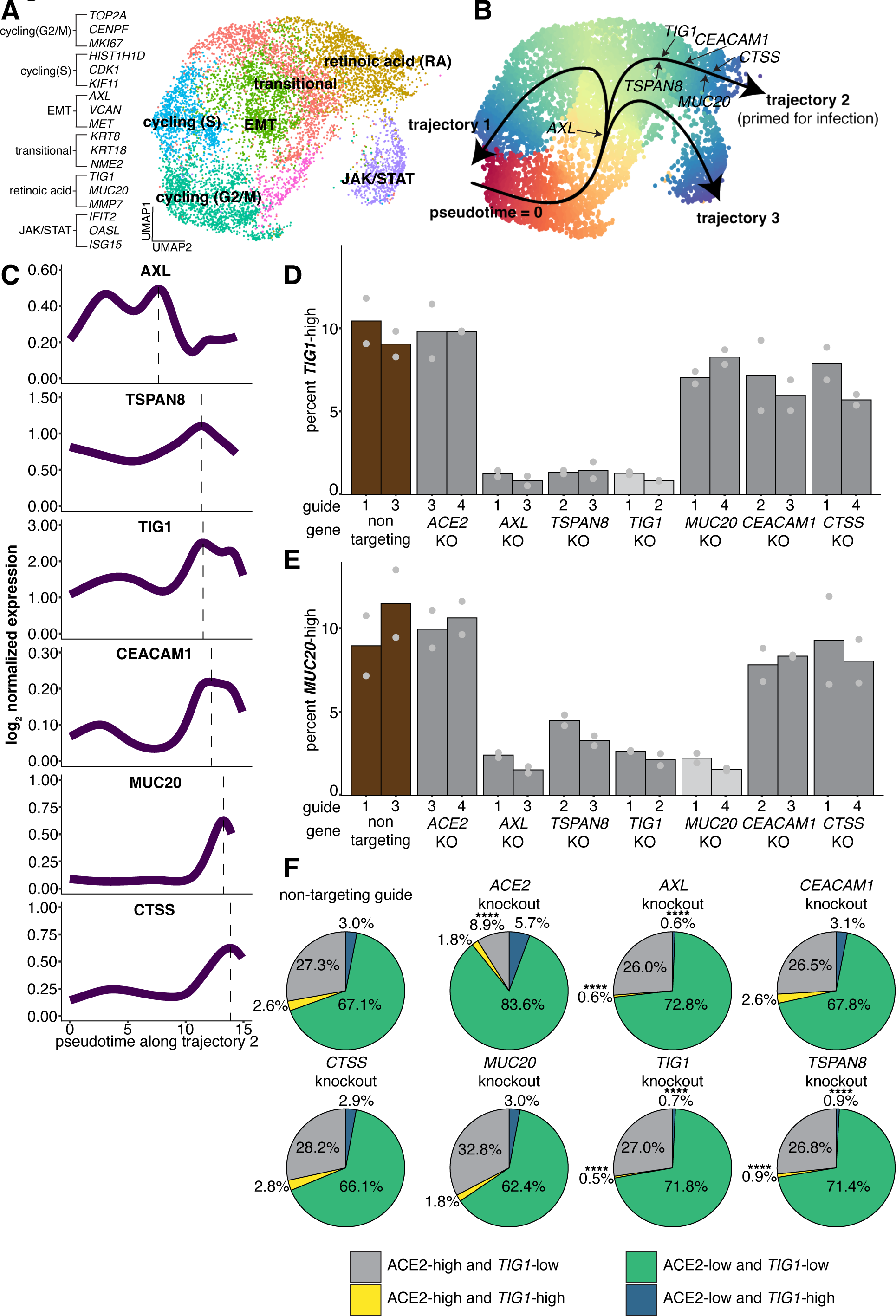
Identification of factors that regulate a cell state that is highly susceptible to infection with SARS-CoV-2. **A.** UMAP representation of single cell RNA-sequencing data collected from Calu-3 cells. **B.** Trajectory analysis performed on Calu-3. Arrow directionality indicates pseudotime ordering along each trajectory. **C.** Normalized expression of factors that promote infection plotted against pseudotime along the *TIG1*-high state trajectory. **D.** single molecule RNA FISH of *TIG1* expression in cells depleted for factors associated with infection. We considered any cell above the 90th percentile of spot counts in the non-targeting control to be *TIG1*-high, and applied that cutoff threshold to all conditions. The top two performing guides from previous experiments were used. Each bar is the average across two biological replicates. **E.** single molecule RNA FISH of *MUC20* expression in the same knockout cells. We considered any cell above the 90th percentile of spot counts in the non-targeting control to be *MUC20*-high, and applied that cutoff threshold to all conditions. The top two performing guides from previous experiments were used. Each bar is the average across two biological replicates. **F.** Co-staining for ACE2 protein and single molecule RNA FISH against *TIG1* in each knockout condition. Significance was assigned by performing a z-test to compare the proportion of each population within each knockout to the proportion of each population in the non-targeting control. Four stars represent a p-value < 0.01 after correcting for multiple comparisons.

We then asked where along the *TIG1*-high trajectory (trajectory 2) each factor identified as important for infection falls. We found that *AXL*, *TSPAN8*, and *TIG1* expression precedes that of *CEACAM1*, *MUC20*, and *CTSS* in pseudotime, with the latter present towards the end of the trajectory (Figure 4C). Such a result suggests that *AXL*, *TSPAN8,* and *TIG1* may be regulating a cell state in which *CEACAM1, MUC20,* and *CTSS* are expressed.

The suggested expression ordering along trajectory 2 is *AXL, TSPAN8, TIG1,* then *CEACAM1, MUC20,* and *CTSS*. To test whether this temporal ordering reflected an underlying regulatory ordering, we knocked out each factor and measured the expression of *TIG1* and *MUC20* to see which factors were potentially upstream regulators of their expression. We found that knocking out *AXL* decreased the percentage of cells with high expression of *TIG1* from 9.89% to 1.01% and *MUC20* from 9.95% to 1.95%. Likewise, knocking out *TSPAN8* also decreased the percentages of cells with high expression of *TIG1* from 9.89% to 1.92% and *MUC20* from 9.95% to 4.13% (Figure 4D-E, Figure S8A-B). *TIG1* knockout also led to a decrease in *MUC20* expression (from 9.95% to 2.36%), while knocking out MUC20 only impacted *MUC20* and not *TIG1* (Figure 4D-E, Figure S8A-B).

The result that *AXL* and *TSPAN8* regulate *TIG1* and *MUC20* suggests that they are upstream of both *TIG1* and *MUC20* in the regulatory chain. Further, the finding that knocking out *TIG1* affects *MUC20* levels but not the other way around indicates that *MUC20* is downstream of *TIG1*. The fact that knocking out *AXL and TSPAN8* affected both *TIG1* and *MUC20* suggests that *AXL* and *TSPAN8* are affecting a broader cellular state rather than just the regulation of expression of one particular gene. Meanwhile, knocking out the factors *CEACAM1* and *CTSS* did not affect *TIG1* or *MUC20* mRNA levels, suggesting that they are terminal factors expressed at the end of the trajectory. Together, these results validate the ordering suggested by our pseudotime analysis.

Our results suggest a regulatory cascade in which *AXL* and *TSPAN8* are upstream of *TIG1*, which in turn is upstream of *MUC20*. *AXL* and *TSPAN8* have been previously reported to directly impact viral entry (Bohan et al., 2021; Hysenaj et al., 2021; Wang et al., 2021), but their regulation of *TIG1* and *MUC20* as well as their known roles in EMT (Antony & Huang, 2017; Vuoriluoto et al., 2011) and epithelial cell differentiation (Zhu et al., 2019) raise the possibility that they can also affect viral infection by driving cells into a more permissive state. *MUC20* behaves similarly to that of ACE2 across assays, with a strong effect in the entry assay but minimal effect on *TIG1* mRNA levels. As *MUC20* knockout had a minimal effect on ACE2 protein levels (Figure 4F), this decrease in viral entry appears to be independent of ACE2 and therefore may point to a more direct role for *MUC20* in promoting viral entry.

### Primed state regulators control only a subset of ACE2 expressing cells

We found that *TIG1* is a marker for a subset of ACE2 expressing cells that are enriched for infection. We also observed a minor relative decrease in ACE2 protein when *AXL*, *TSPAN8*, *TIG1,* and *MUC20* were knocked out (Figure 3F), which could be because these knockouts specifically reduce the small percentage of ACE2- and *TIG1*-high cells in the population. To determine if this decrease in ACE2 was due to a specific depletion of ACE2 expressing cells that are also *TIG1-*high, we co-labeled cells using *TIG1* single molecule RNA FISH and ACE2 immunofluorescence that were depleted of each gene of interest. We found that knockout of *AXL* and *TSPAN8* specifically depleted ACE2-high cells that were also *TIG1*-high (from 2.6% in the non-targeting guide to 0.6% in the case of *AXL*, 0.9% in the case of *TSPAN8*) (Figure 4F). We observed similar trends when *TIG1* was depleted, validating our knockout assay. We found no difference in the percentage of ACE2-high/*TIG1*-low cells within these knockouts (27.3% in the case of the non targeting guide, and 26.0% in the case of *AXL*, 26.8% in the case of *TSPAN8*, and 27.0% in the case of *TIG1*) (Figure 4F). Of note, we found that in the ACE2 knockout condition, the relative decrease in the proportion of ACE2-high and *TIG1*-high cells compared to the non-targeting control (from 2.6% to 1.8%, a reduction of 30.8%) was less than that of the decrease in the proportion of ACE2-high and *TIG1*-low cells (from 27.3% to 8.9%, a reduction of 67.4%) (Figure 4F), perhaps due to differences in ACE2 protein levels amongst the starting populations (Figure S8C-D). Our findings indicate that the factors identified as regulators of *TIG1* expression led to the depletion of a subset of ACE2-high cells that were also *TIG1-*high rather than just a loss of *TIG1* expression, providing additional evidence that these factors may regulate a cell state that is predisposed to infection in addition to their other more direct role in viral entry.

### TIG1-high cells are present within in vivo lung

Having identified a transcriptional state that primes the cell for infection with SARS-CoV-2, we next explored whether a similar state could be found in the human respiratory tract. To do so, we used publicly available single cell RNA sequencing data from human lungs (Habermann et al., 2020). We first filtered the data to only include epithelial cells (*EPCAM*-high/*CDH1*-high), as these are the cell types that are the main targets of SARS-CoV-2 infection and are the most related to Calu-3 cells (Ravindra et al., 2021). After correcting for differences between patients (Korsunsky et al., 2019) (see methods), we identified the known cell types within the lung epithelium: basal cells (*KRT5*-high) (Morrisey, 2018), type I pneumocytes (*AGER*-high) (Acheampong et al., 2022), type II pneumocytes (*SFTPC*-high) (Yao et al., 2021), club cells (*SCGB1A1*-high) (Rawlins et al., 2009), and ciliated cells (*SNTN*-high) (Konishi et al., 2016) (Figure 5A-B, Figure S9A). Next, we used this cell type map to query the expression status of the markers that we identified as important for SARS-CoV-2 infection. *AXL* is expressed in basal stem cells while *TSPAN8* is expressed in a subset of basal stem cells as well as club cells, supporting both factors’ potential roles as drivers of a precursor-like phenotype in Calu-3 cells (Figure 5C). Both *TIG1* and *MUC20* are expressed in a subset of club and ciliated cells, cell types that are infected by SARS-CoV-2 in the lung (Melms et al., 2021; Weber, 2021) (Figure 5C). To confirm that the *TIG1*-high state and not just discrete expression of *TIG1* and *MUC20* was present *in vivo*, we compared the top 100 differentially expressed genes identified in the *TIG1*-high Calu-3 cluster to those identified in *TIG1*-high *in vivo* lung epithelial cells. We found significantly more shared genes than expected by random sampling across the two (20 genes in the case of *TIG1*-high Calu-3 compared to an average of 3.2 in the random sampling case over n=1000 iterations; p-value=0.001), including *CEACAM1, MMP7, MDK, SLPI, AGR2,* and *HLA-DRB1* (Figure S10A-C). Thus, we found that the *TIG1*-high state is similar between cultured cells and *in vivo* lung cells, raising the possibility that this state may reveal the cells most susceptible to infection *in vivo*.

**Figure 5:**
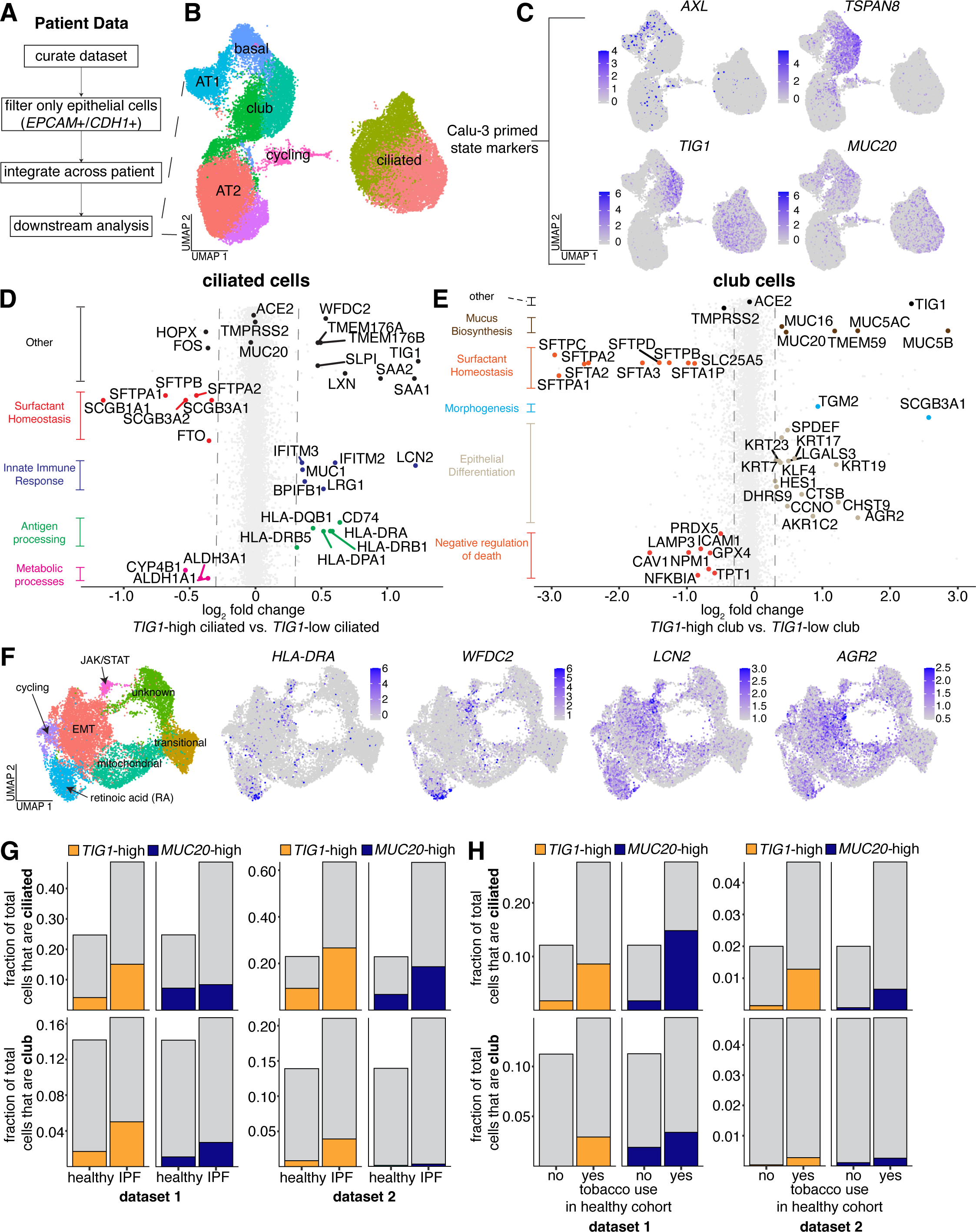
Heterogeneous expression of the *TIG1*-high state is found in distinct cell types in vivo. **A.** Workflow for analysis of public single cell RNA sequencing data from human lungs. Starting from raw count tables, we first performed dimensionality reduction and clustering to identify epithelial cells. We then subset the data to only include clusters that contained epithelial marker gene expression. After subsetting, we re-performed all normalization, variable feature identification, and dimensionality reduction steps. **B.** UMAP of integrated data across patients. Cell types were annotated based on canonical markers of each type (AT1; *AGER*, AT2; *SFTPC*, club; *SCGB1A1*, ciliated; *SNTN*, basal; *KRT5*, cycling; *MKI67*). **C.** Normalized *in vivo* expression of markers associated with Calu-3 that are primed for infection. **D.** To determine genes that are differentially expressed between *TIG1*-high and *TIG1*-low ciliated cells, we first used normalized counts to bin ciliated cells as either *TIG1*-high or *TIG1*-low based on *TIG1* expression. After annotating cells, we ran DESeq2 to identify genes that are both up and down regulated across the bins. To determine significance, we used a p-value cutoff < 0.01. To approximate the biological function of each subset of ciliated cells, we performed GO analysis on the list of genes that were either up or down regulated. **E.** We performed the same analysis as **D,** but instead looked for differentially expressed genes between *TIG1*-high and *TIG1*-low club cells. **F.** Mapping of select genes that are upregulated in **D** and **E** onto the original single cell RNA sequencing dataset obtained from Calu-3 cells. **G.** Fraction of the total cells in healthy or IPF lungs that are ciliated or club. Of that fraction, we then calculated the proportion that was high for *TIG1* or *MUC20* expression. We performed this analysis on two independent publicly available datasets. For dataset 1; n=9 patients in the healthy cohort; n=12 patients in the IPF cohort. For dataset 2; n=28 patients in the healthy cohort; n=32 patients in the IPF cohort. **H.** Fraction of the total cells in the lungs of tobacco users or not tobacco users that are ciliated or club. Of that fraction, we then calculated the proportion that was high for *TIG1* or *MUC20* expression. We performed this analysis on two independent publicly available datasets. For dataset 1; n=7 patients in the tobacco cohort; n=2 patients in the no tobacco cohort. For dataset 2; n=2 patients in the tobacco cohort; n=6 patients in the no tobacco cohort.

Since both ciliated and club cells can be infected with SARS-CoV-2 and both express *TIG1* at variable levels, we next wondered how ciliated and club cells in a *TIG1*-high state differ transcriptionally from the bulk population of each type. We conducted differential expression analysis within both ciliated and club cells, generating two bins for each type depending on a given cell’s *TIG1* expression level. In both comparisons, we saw no difference in the expression of *ACE2*. Across ciliated cells, there was no difference in *TMPRSS2* expression, but *TIG1*-high club cells had lower expression of *TMPRSS2* relative to *TIG1-*low club cells (Figure 5D-E). Our results suggest that *TIG1*-high ciliated cells have higher expression of antigen processing pathways (*HLA-DRB1, HLA-DRB5, HLA-DRA, CD74)* and innate immune-related genes (*IFITM2, IFITM3)*, as well as lower expression of club cell markers and surfactants *(SCGB3A1, SCGB3A2, SFTPB, SFTAP1)* (Figure 5D).

When we performed a similar analysis on *TIG1*-high club cells, we found higher expression of mucus synthesis (*MUC20, MUC5B, MUC5AC, MUC16)* and epithelial differentiation markers (*KRT7, KRT17, KRT23, DHRS9, SCGB3A1, AGR2)*, along with a decrease in expression of type 2 alveolar cell markers (*SFTPA1, SFTPA2, SFTPC, SFTPB)* (Figure 5E). The above results point to the *TIG1*-high state marking subsets of highly differentiated cells with unique phenotypic roles within both ciliated and club cell populations.

Having identified genes that mark *TIG1*-high ciliated and *TIG1*-high club cells *in vivo*, we next mapped genes upregulated within these populations back to our initial Calu-3 single cell RNA sequencing dataset. We found that some *TIG1-*high ciliated cell markers, such as *HLA-DRA* and *WFDC2*, were also highly expressed in *TIG1*-high Calu-3 (Figure 5F). Other genes, such as *LCN2* and *AGR2,* were also present in *TIG1*-high cells, but were expressed more broadly across Calu-3 subtypes (Figure 5F). These data suggest that the *TIG1*-high population identified as the most susceptible to infection in Calu-3 may share transcriptional similarities with *TIG1*-high ciliated and club cells found *in vivo*.

### *TIG1* expression varies across individuals and disease

The severity of illness following SARS-CoV-2 infection varies widely between individuals. Some comorbidities, such as hypertension, diabetes, and chronic inflammatory diseases may also increase the risk of severe infection in certain populations (Ng et al., 2021; Yang et al., 2020). One possibility is that individuals with increased risk of severe illness may have more cells of the *TIG1*-high subtype, promoting infection. To determine if higher levels of *TIG1* expression are present in diseased lungs, we analyzed single cell transcriptome datasets of lung epithelia from patients in various disease cohorts linked to fibrosis and inflammation (Habermann et al., 2020). We found an increase in the relative proportion of ciliated cells within patients with idiopathic pulmonary fibrosis (IPF) compared to the healthy group (Figure S9B, Figure S11B). In addition to an absolute increase in the number of ciliated cells, we saw an increase in the frequency of *TIG1-*high and *MUC20*-high expressing ciliated and club cells in individuals with IPF (Figure 5G). While we found variability in *TIG1* and *MUC20* expression between patients in each disease group, such variability was not large enough to explain the absolute differences we observed in the fraction of high expressing cells for each gene (Figure S10D-E). We confirmed these results by analyzing a second public dataset in the same manner as above (Adams et al., 2020) (Figure 5G, Figure S11A-B). We also found an increase in the frequency of *TIG1-*high ciliated cells in individuals with chronic hypersensitivity pulmonitis (cHP) (Figure S9C). These data suggest that certain inflammatory diseases have more cells expressing *TIG1* (and therefore, potentially in the primed *TIG1*-high state) compared to healthy lungs. We further found patients diagnosed with cHP, non-specific interstitial pneumonia (NSIP), and interstitial lung disease (ILD) enriched for club cells, but not ciliated cells, expressing high levels of *MUC20* (Figure S9C), suggesting that *MUC20* and *TIG1* may be enriched in different cell types of the lung in a disease-specific manner.

We also found variability in the expression of *TIG1* and *MUC20* across patients in the healthy cohort (Figure S9E), raising the possibility that the expression of these factors might be related to the variability in disease outcomes observed across healthy patients infected with SARS-CoV-2. We found no correlation between the expression of these genes and patient age (Figure S9F), but did find an increase in expression of both markers in ciliated and club cells within individuals who used tobacco compared to those that did not (Figure 5H). We confirmed this increase by analyzing an additional dataset (Figure 5H, Figure S12A-B) (Reyfman et al., 2019). Altogether, these analyses suggest the possibility that the proportion of cells in the *TIG1*-high state can vary across individuals and may be affected by environment or disease, potentially driving variable SARS-CoV-2 infection outcomes across different populations.

## Discussion

Viral tropism impacts susceptibility to infection across multiple biological scales, from species to organisms within the species to cell types within the organism. At the molecular level, tropism is often ascribed to the presence or absence of a specific group of factors known to be required by the virus in a particular cell type. However, we know that the presence of these canonical factors alone is insufficient to fully explain the variations observed in infection levels and outcomes. We propose that distinct cell states within a specific cell type may contribute to differential susceptibility to infection. Identifying the factors specific to these states could be an important means to define tropism with single cell resolution, potentially revealing new host factors necessary for infection. Until now, finding these states has been difficult because the host cells change so rapidly and markedly after infection.

Here, using retrospective clone tracing in Calu-3 cells (Emert et al., 2021), we show that there is a specific cellular state that is highly susceptible to infection with SARS-CoV-2. These data complement other studies that have relied on traditional single cell RNA sequencing methods (Fiege et al., 2021; Melms et al., 2021; Ravindra et al., 2021). The factors expressed within this state include previously described factors, as well as previously undescribed proteins. This transcriptional signature was also found *in vivo* and was heterogeneously expressed across patients, providing a potential mechanism to explain the variability in SARS-CoV-2 permissivity observed across individuals. Moreover, this detailed transcriptional state provides a more precise description of the specific subset of cells within the human respiratory tract that may exhibit heightened susceptibility to infection. These cells potentially play a pivotal role in the early stages of lung infection, where cells are infected under limiting conditions.

Our approach provides an important complement to genetic screens in the identification of host factors important for the viral life cycle (Baggen et al., 2021; Bailey & Diamond, 2021; Biering et al., 2022; Israeli et al., 2022; Rebendenne et al., 2022; Wei et al., 2021; Zhu et al., 2021). Rewind can provide information about the entire cellular state. It may also reveal factors that may not be detectable by screening. A high percentage of the individual factors we identified did validate as important for viral infection, suggesting that profiling cells prior to infection captures a more informed snapshot of the factors that determine whether a cell is highly susceptible to infection with SARS-CoV-2.

In addition to revealing host factors that play direct roles in infection, our work also revealed a new cellular state that promotes infection. We further defined a regulatory trajectory that is associated with this highly infectable state. Exploring whether other respiratory viruses rely on these programs or alternative programs for infection will provide a more general overview of viral infection in the lung. In addition, the lung epithelium shares phenotypic similarities to other systems, such as the gut, perhaps suggesting that similar programs may drive variable susceptibility of other cell types to viruses outside of the lung.

Future work will determine the mechanism by which the *TIG1-*high state is preferentially infected. One possibility is that cells in a *TIG1-*high state are *enriched* for specific factors that are pro-viral. Indeed, we found that factors expressed in this state, such as AXL, TSPAN8, TIG1, and MUC20, do promote infection. However, we also identified a state that is characterized by the expression of innate immune factors that appears to be resistant to infection. Further characterization of this state and its molecular drivers may reveal new strategies for promoting resistance to infection. Determining the full spectrum of cellular states within a population of cells exposed to virus not only has the potential to provide a more detailed picture into the mechanisms that drive infection, but may also help formulate novel antiviral strategies.

## Methods

### Cell lines and culture

Unless otherwise noted, we cultured all cells at 37°C and 5% CO_2_. We expanded Calu-3 (HTB-55) cells in growth medium on TC plastic dishes, and split cells 1:5 upon reaching 80% confluency. For general maintenance, we used a growth medium containing MEM Earles with Glutamax (11095114) + 10% FBS (16000044) + 1X Glutamax (35050061) and 0.5ug/mL puromycin. When passaging cells, we performed dissociation with TrypLE (12604021) by incubation at 37°C for 10 minutes.

### Virus production

Viral stocks were prepared as previously described (Schultz et al., 2022). SARS-CoV-2 WA1 was provided by the BEI/CDC. Virus stocks were amplified using the ARTIC primer set and sequenced using the MinION system (Oxford Nanopore Technologies) by the J. Craig Venter Institute (MD, USA) to more than 4,000× genome coverage. Stock sequence was verified by aligning reads to the reference genome provided by the BEI (GSAID accession: EPI_ISL_890360) using minimap2 version 2.22 with the ‘map-ont’ presets, followed by inspection of the consensus sequence and alignment using IGV. Stocks had less than 1% variation. Stock virus was prepared by infection of Vero E6 cells expressing TMPRSS2 in growth medium (DMEM (Quality Biological), supplemented with 10% (v/v) fetal bovine serum (Gibco), 1% (v/v) penicillin–streptomycin (Gemini Bio-products) and 1% (v/v) L-glutamine (2 mM final concentration; Gibco) fetal bovine serum plus for 2 or 3 days when cytopathic effect (CPE) was visible. Media were collected and clarified by centrifugation before being aliquoted for storage at −80 °C. Titre of stock was determined by plaque assay or 50% tissue culture infectious dose (TCID50) analysis using Vero E6 cells as previously described (Schultz et al., 2022). All work with infectious virus was performed in a biosafety level 3 laboratory and approved by the University of Pennsylvania Biosafety Committee.

### RT-qPCR

Calu-3 cells (150,000 cells per well) were plated in 24-well plates. One day later, cells were infected with SARS-CoV-2 (MOI = 0.3) for 1 hour and, subsequently, the virus inoculum was removed. The cells were placed into fresh medium daily with the indicated drugs. Total RNA was purified using TRIzol (Invitrogen) followed by the RNA Clean and Concentrate kit (Zymo Research) 48 h after infection for Calu-3. For cDNA synthesis, reverse transcription was performed with random hexamers and Moloney murine leukaemia virus (M-MLV) reverse transcriptase (Invitrogen). Synthesized RNA was used as a standard (BEI). Gene-specific primers to SARS-CoV-2 (Wuhan v1, NSP14) and SYBR green master mix (Applied Biosystems) were used to amplify viral RNA, and 18S rRNA primers were used to amplify cellular RNA using the QuantStudio 6 Flex RT–PCR system (Applied Biosystems). Relative quantities of viral and cellular RNA were calculated using the standard curve method. Viral RNA was normalized to 18S RNA for each sample (Wuhan V1/18S).

### dsRNA immunofluorescence

Cells were infected with SARS-CoV-2 (multiplicity of infection (MOI) = 0.5). Cells were fixed 40–48 h post-infection in 4% formaldehyde in PBS for 15 min at room temperature, washed three times with PBS, blocked with 2% BSA in PBST for 60 min, and incubated in primary antibody (anti-double-stranded RNA J2, absolute antibody, 1:500) overnight at 4 °C. Cells were washed three times in PBST with an automated plate washer (BioTek) and incubated in secondary antibody (anti-mouse Alexa 488, 1:1,000 and Hoescht 33342) for 1 h at room temperature. Cells were washed three times in PBST with an automated plate washer and imaged using an automated microscope (ImageXpress Micro, Molecular Devices). Cells were imaged with a ×10 objective, and four sites per well were captured. The total number of cells and the number of infected (double-stranded RNA+) cells were measured using the cell scoring module (MetaXpress 5.3.3), and the percentage of infected cells was calculated.

### VSV infections

Calu-3 cells (30,000 cells per well) were plated in 96-well plates. One day later, cells were infected with either VSV-GFP (WT VSV glycoprotein, MOI 162) or VSV-GFP-Spike (with SARS-CoV-2 spike protein, MOI 25). One day post infection, cells were fixed with 4% PFA solution and subsequently imaged for viral infection quantifying GFP positivity and nuclei. VSV-GFP was grown and titered in BHK cells and VSV-GFP Spike was grown and titered in Vero CCL81 cells.

### Clone barcode lentivirus library generation

Barcode libraries were constructed as previously described (Emert et al., 2021; Goyal et al., 2021; Jiang et al., 2022). The full protocol is available at https://www.protocols.io/view/barcode-plasmid-library-cloning-4hggt3w. Briefly, we modified the LRG2.1T plasmid, kindly provided by Junwei Shi, by removing the U6 promoter and single guide RNA scaffold and inserting a spacer sequence flanked by EcoRV restriction sites just after the stop codon of GFP. We digested this vector backbone with EcoRV (NEB) and gel purified the resulting linearized vector. We ordered PAGE-purified ultramer oligonucleotides (IDT) containing 30 nucleotides homologous to the vector insertion site surrounding 100 nucleotides with a repeating “WSN” pattern (W = A or T, S = G or C, N = any) and used Gibson assembly followed by column purification to combine the linearized vector and barcode oligo insert. We performed 9 electroporations in total of the column-purified plasmid into Endura electrocompetent *E. coli* cells (Lucigen #60242-1) using a Gene Pulser Xcell (Bio-Rad #1652662), allowing for recovery before plating serial dilutions and seeding cultures (200 mL each) for maxipreparation.

We incubated these cultures on a shaker at 225 rpm and 32 °C for 12–14 h, after which we pelleted cultures by centrifugation and used the EndoFree Plasmid Maxi Kit (Qiagen #12362) to isolate plasmid according to the manufacturer’s protocol. Barcode insertion was verified by polymerase chain reaction (PCR) from colonies from plated serial dilutions. We pooled the plasmids from the separate cultures in equal amounts by weight before packaging into lentivirus. We estimated our library complexity as described elsewhere (Goyal et al., 2021). Briefly, we sequenced three independent transductions in WM989 A6-G3 melanoma cells and took note of the total and pairwise overlapping extracted barcodes. Using the mark-recapture analysis formula, we estimate our barcode diversity from these three transductions to be between 48.9 and 63.3 million barcodes.

### Lentivirus packaging of clone barcode library

Before plasmid transfection, we first grew HEK293FT to near confluency (80-95%) in 10cm plates in DMEM + 10% FBS without antibiotics. For each 10cm plate, we added 80 µL of polyethylenimine (Polysciences #23966) to 500 µL of Opti-MEM (Fisher #31985062) and separately combined 5 µg of VSVG, 7.5 µg of pPAX2, and 7.35 µg of the barcode plasmid library in 500 µL of Opti-MEM. We mixed both solutions, vortexed for 20 seconds, and incubated the combined plasmid-polyethylenimine solution at room temperature for 15 min. We added 1.09 mL of the combined plasmid-polyethylenimine solution dropwise to each 10cm dish. After 4-6 hours, we aspirated the transfection media from the HEK293FTs and added Calu-3 medium containing MEM + 10% FBS + 1x Glutamax. After 48 hours, filtered the virus-laden media through a 0.45µm PES filter (Millipore-Sigma #SE1M003M00) and stored 1 ml aliquots in cryovials at -80°C.

### Transduction of Calu-3 with lentiviral clone barcodes

To transduce Calu-3 cells with our lentiviral barcode library, we freshly thawed virus-laden media on ice, and added 100uL of viral stock + 4ug/mL of polybrene (Millipore-Sigma #TR-1003-G) to Calu-3 media containing 100,000 live cells. Cells were then plated into one well of a 6-well plate. The volume of viral stock used was decided by measuring the multiplicity of infection (MOI) with different viral titers. For single-cell RNA-sequencing experiments, we aimed for a low MOI with ∼10%-25% to ensure at most one barcode integration event per cell. After ∼8 hours, viral laden media was removed from the cells and replaced with standard Calu-3 media. Prior to single cell RNA sequencing experiments, the barcoded cells (GFP-positive) were sorted to recover only cells that received a barcode. Unless specified, barcoded cells were passaged for ∼3 population (∼5-7 days) doublings before downstream assays were performed.

### SARS-CoV-2 single molecule RNA FISH probe design

To target the SARS-CoV-2 genome, we referenced the ORF1a sequence of the first US isolate of SARS-CoV-2 (USA-WA1/2020) from NCBI (Genbank MN985325.1). We designed complementary oligonucleotide probe sets using custom probe design software (MATLAB). For each probe, we then performed a local blast against the human transcriptome and Nucleic Acids of Coronavirus and other Human Oronasopharynx pathogens (NACHO), a database we created of 562,446 sequences of other viruses that infect the human respiratory tract. All probes in the top 5% of hits based on E-value and bit score were excluded. We then ordered probes with a primary amine group on the 3′ end from Biosearch Technologies (Table S2 for probe sequences). We then pooled all oligonucleotides and coupled the set to Cy3 (GE Healthcare), Alexa Fluor 594 (Life Technologies) or Atto647N (ATTO-TEC) N-hydroxysuccinimide ester dyes.

### single molecule RNA FISH in suspension and FACS sorting of fixed infected cells

In order to separate infected cells from not infected cells for barcode recovery, we used a modified version of our previously described single molecule RNA FISH protocol (Raj et al., 2008) to label cells containing viral RNA. Briefly, 48 hours post infection, cells were washed, dissociated, and fixed in suspension with 4% PFA solution for 15 minutes. Following fixation, the pellet was washed 3x with 1xDPBS and stored in 70% EtOH overnight at 4°C. For hybridization of viral RNA FISH probes in suspension, we washed samples with wash buffer (10% formamide in 2X SSC) before adding hybridization buffer (10% formamide and 10% low molecular weight dextran sulfate in 2X SSC) with 100 ng of single molecule RNA FISH probe per one million cells in one mL of hybridization buffer. Samples were incubated overnight at 37°C. 12-18 hours later, we performed two 30-min washes at 37°C with the wash buffer, after which we added 2X SSCT with 50 ng/mL of DAPI for flow sorting. Immediately following single molecule RNA FISH, FACS (FACS Aria FUSION) was performed on FISH signal and cells were binned into infected and not infected groups based on signal intensity. Each bin was collected into 1mL DNA lo-bind Eppendorf tubes (part number 0030122275) containing 400 μL of 2xSSCT.

### Sequencing clone barcodes from genomic DNA

We prepared barcode libraries from genomic DNA (gDNA) with a modified protocol from previously described work (Emert et al., 2021; Goyal et al., 2021; Jiang et al., 2022). Briefly, we isolated gDNA from fixed barcoded cells using the QIAmp DNA FFPE Kit (Qiagen #56404) per the manufacturer’s protocol. We then immediately performed targeted amplification of the barcode using custom primers containing Illumina adaptor sequences, unique sample indices, variable-length staggered bases, and an “UMI” consisting of 6 random nucleotides (NHNNNN). We determined the number of amplification cycles (N) by initially performing a separate quantitative PCR (qPCR) and selecting the number of cycles needed to achieve one-third of the maximum fluorescence intensity for serial dilutions of genomic DNA. The thermal cycler (Veriti #4375786) was set to the following settings: 98°C for 30 sec, followed by N cycles of 98°C for 10 sec and then 65°C for 40 sec and, finally, 65 °C for 5 min. Upon completion of the PCR reaction, we immediately performed a 0.7X bead purification (Beckman Coulter SPRISelect #B23319), followed by final elution in warm nuclease-free water. Purified libraries were quantified with a High Sensitivity dsDNA kit (Thermo Fisher #Q33230) and on a Qubit Fluorometer (Thermo Fisher #Q33238), pooled, and sequenced on a NextSeq 500 using 150 cycles for read 1 and 8 reads for each index (i5 and i7). The primers used are described in (Goyal et al., 2021).

### Analysis of sequenced barcodes from genomic DNA

Barcode libraries amplified from genomic DNA-sequencing data were analyzed as previously described (Emert et al., 2021), with the custom barcode analysis pipeline available at https://github.com/arjunrajlaboratory/timemachine. This pipeline searches for a barcode sequence that satisfies a minimum phred score, a minimum length, and correct flank sequences. After extracting barcode sequences, we use STARCODE (Zorita et al., 2015), available at https://github.com/gui11aume/starcode, to merge sequences with Levenshtein distance ≤ 8 and add the counts across collapsed (merged) barcode sequences and samples. We then used a median based normalization to correct for sequencing depth across samples.

### Simulation for barcode overlap

We adapted a computational model developed previously (Jain et al., 2023) to simulate the barcode overlap expected in our experiments. Briefly, this model simulates all steps of our experimental design, including material loss and cell death. We performed 1000 independent simulations to obtain a distribution of barcode overlap values that were then used to assign significance to the observed overlap value obtained in our experiments.

### Single-cell RNA-sequencing

We used the 10X Genomics single-cell RNA-seq kit v3 to sequence barcoded cells. We resuspended the cells (aiming for up to ∼12,000 cells for recovery/ sample) in PBS and followed the protocol for the Chromium Next GEM Single Cell 3ʹ Reagent Kits v3.1 as per manufacturer directions (10X Genomics, Pleasanton, CA). Briefly, we generated gel beads-in-emulsion (GEMs) using the 10X Chromium system, and subsequently extracted and amplified barcoded cDNA as per post-GEM RT-cleanup instructions. We then used a fraction of this amplified cDNA (25%) and proceeded with fragmentation, end-repair, poly A-tailing, adapter ligation, and 10X sample indexing per the manufacturer’s protocol. We quantified libraries using the High Sensitivity dsDNA kit (Thermo Fisher #Q32854) and Bioanalyzer 2100 (Agilent #G2939BA). Concentration of libraries were confirmed on a Qubit 2.0 Fluorometer (Thermo Fisher #Q32866) prior to sequencing on a NextSeq 500 machine (Illumina) using 28 cycles for read 1, 55 cycles for read 2, and 8 cycles for i7 index.

### Computational analyses of single-cell RNA-sequencing and expression data

We adapted the cellranger v3.0.2 by 10X Genomics into our custom pipeline (https://github.com/arjunrajlaboratory/10XCellranger) to map and align the reads from NextSeq sequencing run(s). Briefly, we downloaded the bcl counts and used cellranger mkfastq to demultiplex raw base call files into library-specific FASTQ files. We aligned the FASTQ files to the hg38 human reference genome and extracted gene expression count matrices using cellranger count, while also filtering and correcting cell identifiers and unique molecular identifiers (UMI) with default settings. We then performed all downstream analysis within Seurat v3. Within each sample, we removed genes that were present in less than three cells in addition to any cell that had less than or equal to 200 detected genes. We also set a cutoff for mitochondrial gene fraction based on the distribution of fractions across all cells in each experiment. For samples that were technical replicates of one another, we integrated using SCTransform (Hafemeister and Satija, 2019) in accordance with the Satija lab workflow(https://satijalab.org/seurat/articles/integration_introduction.html).

For samples that were treated differently, we utilized harmony (Korsunsky et al., 2019) to prevent over-clustering of our data while still removing batch effects across condition and sample. For each experiment, we used these integrated datasets to reduce the dimensions of our data by principal component analysis (PCA) and Uniform Manifold Approximation and Projection (UMAP). For Figure 1, we utilized the first 30 principal components and clustered cells based on a resolution of 0.3. For Figure 4, we utilized the first 15 principal components and clustered cells based on a resolution of 0.5. For Figure 5, we utilized the first 15 principal components and clustered cells based on a resolution of 0.3. Of note, we identified a cluster in Figure 1 as doublets; marked by feature counts and RNA counts more than four times that of the next nearest cluster. We therefore decided to remove this cluster from downstream analyses. To calculate the markers for each cluster in UMAP space, we used Seurat v3s FindAllMarkers function with the default settings. To calculate differentially expressed genes based on expression rather than cluster, we first computationally sorted cells based on expression of a marker of interest. We then used the zinbwave function (Risso et al., 2018) in R to generate a zero-inflated negative binomial model to represent our data in low dimensions. We used the weights of this model directly in DESeq2 (Love et al., 2014) to analyze differentially expressed genes using the following parameters; test = "LRT", sfType = "poscount", reduced = ∼1, useT=TRUE, minmu=1e-6. For GO analysis, we loaded differentially expressed genes into the shiny app located at http://bioinformatics.sdstate.edu/go/. To assign gene expression signature scores, we used Seurat v3s AssignModuleScore function with default settings for a list of genes of interest.

### Barcode recovery from single-cell RNA-sequencing data

We extracted barcode information directly from the amplified cDNA from 10X Genomics V3 chemistry protocol (step 2). We ran a PCR side reaction with one primer that targets the 3’ UTR of GFP and the other that targets a region introduced by the amplification step within the V3 chemistry of 10X genomics (“Read 1”). The two primers amplify both the 10X cell-identifying sequence as well as the 100 bp barcode that we introduced lentivirally. The number of cycles, usually between 10-12, are decided by the Ct value from a qPCR reaction (NEBNext Q5 Hot Start HiFi PCR Master Mix (New England Biolabs)) for the specified cDNA concentration. The thermal cycler (Veriti 4375305) was set to the following settings: 98 °C for 30 s, followed by N cycles of 98 °C for 10 s and then 65 °C for 2 min and, finally, 65 °C for 5 min. Upon completion of the PCR reaction, we immediately performed a 0.7X bead purification (Beckman Coulter SPRISelect) followed by final elution in nuclease-free water. Purified libraries were quantified with a High Sensitivity dsDNA kit (Thermo Fisher) on Bioanalyzer 2100 (Agilent #G2939BA), pooled, and sequenced on a NextSeq 500. We sequenced 26 cycles on Read 1 which gives 10X cell-identifying sequence and UMI, 124 cycles for read 2 which gives the barcode sequence, and 8 cycles for index i7 to demultiplex pooled samples. The primers used are described in (Goyal et al., 2021).

### Computational analyses of barcoded single-cell datasets

The barcodes from the side reaction of single-cell cDNA libraries were recovered by developing custom shell, R, and python scripts, which are all available at this link: https://github.com/arjunrajlaboratory/10XBarcodeMatching. Briefly, we scan through each read searching for sequences complementary to the side reaction library preparation primers, filtering out reads that lack the GFP barcode sequence, have too many repeated nucleotides, or do not meet a phred score cutoff. Since small differences in otherwise identical barcodes can be introduced due to sequencing and/or PCR errors, we merged highly similar barcode sequences using STARCODE software (Zorita et al., 2015), available at https://github.com/gui11aume/starcode. For varying lengths of barcodes (30, 40 or 50, see the pipeline guide provided) depending on the initial distribution of Levenshtein distance of non-merged barcodes, we merged sequences with Levenshtein distance ≤ 8, summed the counts, and kept only the most abundant barcode sequence. For downstream analysis, we first filtered out all barcodes that were below a minimum cutoff of unique molecular identifiers (UMI) counts (dependent on sequencing depth of the experiment). Following filtering, we successfully matched a barcode to ∼40-60% of cells in each experiment. In order to determine the significance of barcode enrichment within each cluster in our Rewind experiments, we first generated a starting distribution of barcodes in the experiment by randomly sampling without replacement the same number of barcodes as each sample of interest for the total pool of barcodes recovered (generating hundreds of ‘pseudo-samples’ of same size as the actual samples of interest). Using that random distribution, we determined the baseline fraction of barcodes that one would expect in each cluster in UMAP space for each sample size. In order to assign significance to each cluster, we computed a z-score comparing the actual fraction of barcodes recovered in each cluster for each sample to the sampled distribution. In the case of Figure 1, where we directly compared two samples against each other within each cluster, we took the difference of each sample’s z-score compared to the sampled distribution.

### Trajectory analysis and SCENIC implementation

For pseudotime ordering, we used the slingshot package (Street et al., 2018) to fit trajectories to our single cell data and then the tradeSeq package (Van den Berge et al., 2020) to calculate differentially expressed genes along each trajectory and visualize expression. We followed the in depth workflow listed here: https://www.bioconductor.org/packages/release/bioc/vignettes/tradeSeq/inst/doc/tradeSeq.html. To infer gene regulatory networks, we used the SCENIC package (Aibar et al., 2017) using default parameters.

### HCR in suspension protocol

To perform HCRv3 in suspension, we first designed HCR oligonucleotide probes to target our gene of interest. We used the probe designer described above to design non-overlapping 52-mer oligos with a target Gibbs free energy for binding of −60 (allowable Gibbs free energy [−70, −50]). We then divided each 52-mer oligo into two non-overlapping 25-mer sequences (removing the middle two nucleotides) and appended split-initiator HCR sequences using a custom matlab script (see Supp. Table 1 for probe sequences). We then proceeded with an adapted version of the manufacturer’s protocol. We fixed dissociated cells in suspension by washing the cells with 1xDPBS, resuspending the cell in ice-cold 1xDPBS, adding an equal volume of ice-cold fixation buffer (3.7% formaldehyde 1xPBS) and then incubating with rotation at room temperature for 10 min. We next pelleted fixed cells by centrifugation at 800*g* for 3 min, washed twice with ice-cold 1xPBS and then resuspended in 70% ethanol and stored fixed cells at 4°C. For primary probe hybridization, we used 1.2pmol of probe for each gene of interest in 0.5mL of hybridization buffer (30% formamide, 10% dextran sulfate, 9 mM citric acid pH 6.0, 50 μg ml−1 of heparin, 1xDenhardt’s solution (Life Technologies, 750018) and 0.1% Tween-20 in 5xSSC). Following a 30 minute prehybridization incubation at 37°C, samples were incubated in the probe-hybridization mixture for 6 hours at 37°C. After the primary probe hybridization, we washed samples 4x5 minutes at 37°C with washing buffer (30% formamide, 9 mM citric acid pH 6.0, 50 μg ml−1 of heparin and 0.1% Tween-20 in 5xSSC) and then incubated the samples at room temperature 2x5 minutes with 5x SSCT (5x SSCT + 0.1% Tween-20. For amplification, we first incubated the samples for 30 minutes at room temperature in amplification buffer (10% dextran sulfate and 0.1% Tween-20 in 5xSSC). During this incubation, we snap cooled individual HCR hairpins (Molecular Instruments) by heating to 95°C for 90 s and then immediately transferring to room temperature to cool for 30 minutes. After these 30 minutes, we resuspended and pooled the hairpin in amplification buffer to a final concentration of 6nM each. We added the hairpin solution to the samples and then incubated the samples at room temperature overnight. The next morning, we washed samples 5x5 minutes with 5xSSCT containing 50 ng ml−1 DAPI prior to imaging.

### RNA FISH on cells in plates

We performed single-molecule RNA FISH as previously described (Raj et al., 2008). For the genes used in this study, we designed complementary oligonucleotide probe sets using custom probe design software (MATLAB) and ordered them with a primary amine group on the 3′ end from Biosearch Technologies (Table S2 for probe sequences). We then pooled each gene’s complementary oligos and coupled the set to Cy3 (GE Healthcare), Alexa Fluor 594 (Life Technologies) or Atto647N (ATTO-TEC) N-hydroxysuccinimide ester dyes. The cells were fixed as follows: we aspirated media from the plates containing cells, washed the cells once with 1X DPBS, and then incubated the cells in the fixation buffer (3.7% formaldehyde in 1X DPBS) for 10 min at room temperature. We then aspirated the fixation buffer, washed samples twice with 1X DPBS, and added 70% ethanol before storing samples at 4°C. For hybridization of RNA FISH probes, we rinsed samples with wash buffer (10% formamide in 2X SSC) before adding hybridization buffer (10% formamide and 10% dextran sulfate in 2X SSC) containing 10ng of each RNA FISH probe per 50uL of buffer and incubating samples overnight in humidified containers at 37°C. The next morning, we performed two 30-min washes at 37°C with the wash buffer, after which we added 2X SSC with 50 ng/mL of DAPI. We mounted the sample(s) for imaging in 2X SSC.

### Immunofluorescence for ACE2

Our protocol for labeling cells with an anti-ACE2 antibody was as follows. First, we washed cells with 500uL PBS + 0.05% Tween20 for 5 minutes at room temperature. We then blocked for 30 minutes at room temperature with 250uL of blocking buffer containing 1% recombinant albumin (B9200S) in 1x PBS + 0.05% Tween20. Following blocking, we replaced the blocking buffer with a fresh 250uL of the same blocking buffer containing a 1:1000 dilution of our primary antibody against ACE2 (polyclonal Rabbit-anti-hACE2, Abcam, Cat# Ab15348). Following a 1 hour incubation at room temperature, we did 3x5 minute washes with 500uL of PBS + 0.05% Tween20. Next, we added 250uL of blocking buffer containing a 1:500 of our secondary antibody (goat anti-rabbit, Alexa594, cat# A-11012). Following a 30 minute incubation, we then washed 3x5 minutes with PBS and stored in PBS until imaging or flow.

### Immunofluorescence and single molecule RNA FISH

To perform both single molecule RNA FISH and immunofluorescence on the same sample, we followed our standard single molecule RNA FISH protocol described above. We then subsequently performed immunofluorescence per the above protocol. It is important to use recombinant (or molecular biology grade, RNAse free) albumin for blocking in this protocol, as we found that standard serum and BSA degrades RNA FISH signal rapidly.

### Immunofluorescence and HCR co-staining

To perform both HCR and immunofluorescence on the same sample, we followed the following protocol. We first performed HCR as described above, through the primary hybridization step (stopping before amplification). Then, prior to amplification, we stained cells according to the immunofluorescence protocol above to completion. After completion of all immunofluorescence steps, we then proceeded with the amplification stage of HCR. It is important to note that FISH methods are sensitive to PBS, so it is critical to limit the samples exposure time to PBS. As with single molecule RNA FISH, we found that HCR was sensitive to non-nuclease free blocking agents so it is recommended to use recombinant albumin or alternatives.

### Generation of CRISPR guide constructs

To knockout identified genes of interest, we generated lentiviral CRISPR-Cas9 constructs. We selected 4 guides per gene from a genome-wide database designed using optimized metrics in (Doench et al., 2016). We ordered pairs of complementary forward and reverse single stranded oligonucleotides (IDT) for each guide containing compatible overhang sites for insertion into a lentiCRISPRv2-Puro (Addgene #98290) backbone, which simultaneously encodes Cas9 and an insertable target gDNA. We resuspended each oligo to 25uM in NF-H2O and performed phosphorylation and annealing of each oligo pair by combining the oligos with 1X T4 ligase buffer (NEB #B0202S) and polynucleotide kinase (NEB #M0201S) and incubating in a thermocycler (Veriti #4375305) with the following settings: 37°C for 30 min, 95°C for 5 min, and ramping down to 25°C at 5°C/min. Then, we inserted our annealed oligos into the lentiCRISPRv2-Puro backbone via Golden Gate assembly; we combined 50 ng of backbone with ∼25 ng of annealed oligo along with T4 ligase buffer (NEB #B0202S), T4 ligase (#M0202S), and BsmBI restriction enzyme (NEB #R0739S) and incubated in a thermocycler (Veriti #4375305) with the following settings: 50 cycles of 42°C for 5 min and 16°C for 5 min followed by 65°C for 10 min. To amplify plasmid before lentiviral packaging, we performed heat-shock transformation of Stbl3 *E. coli* at 42°C cells for each guide before plating on LB plates with ampicillin and incubating at 225 rpm and 37°C for 12-16 hours. Individual colonies were picked for each guide and grown out in 5mL liquid cultures of LB with ampicillin before each culture was pelleted by centrifugation. To isolate plasmids, we used the Monarch Plasmid Miniprep Kit (Monarch #T1010L) according to the manufacturer’s protocol. Guide sequences were verified by Sanger sequencing of the isolated plasmid. Next, we packaged sequence-verified plasmids for each guide into lentivirus. We first grew HEK293T to 65%-80% confluency in 6-well plates in DMEM + 10% FBS without antibiotics and on 0.1% gelatin (#ES-006-B). For each individual plasmid, we combined 0.5 ug of pMD2.G, 0.883 ug of pMDLg, 0.333 ug of pRSV-Rev, and 1.333 ug of the plasmid in 200 uL of Opti-MEM. After vortexing, we added 9.09 uL of polyethylenimine (Polysciences #23966) and incubated for 15 min at room temperature before adding the final mixture for each plasmid to a HEK293-containing well of the 6-well plates. After 4-6 hours, we aspirated the media and added fresh Calu-3 media. After 48 hours, we filtered the virus-laden media through a 0.45µm PES filter (Millipore-Sigma #SE1M003M00) and stored 1.5mL aliquots in cryovials at -80°C.

### Transduction of Calu-3 with lentiviral CRISPR constructs

To transduce Calu-3 cells with CRISPR guides, we thawed each virus-laden media on ice, and added 500 uL of stock to 500,000 cells in 1 mL of volume within a well of a 6 well plate. To each well, we also added 4ug/mL of polybrene (Millipore-Sigma #TR-1003-G). After ∼16 hours of incubation at 37°C, we changed media and began selection in 1 ug/mL of puromycin. This selection was continued for 8 days to ensure the death of all cells that did not receive a guide construct, at which time cells were plated for downstream assays.

### Imaging

To image RNA FISH and nuclei signal, we used a Nikon TI-E inverted fluorescence microscope equipped with a SOLA SE U-nIR light engine (Lumencor), a Hamamatsu ORCA-Flash 4.0 V3 sCMOS camera, and 4X Plan-Fluor DL 4XF (Nikon MRH20041/MRH20045), 10X Plan-Fluor 10X/0.30 (Nikon MRH10101) and 60X Plan-Apo λ (MRD01605) objectives. We used the following filter sets to acquire different fluorescence channels: 31000v2 (Chroma) for DAPI, 41028 (Chroma) for Atto 488, SP102v1 (Chroma) for Cy3, 17 SP104v2 (Chroma) for Atto 647N, and a custom filter set for Alexa 594. We tuned the exposure times depending on the dyes used (Cy3, Atto 647N, Alexa 594, and DAPI). For large scans, we used a Nikon Perfect Focus system to maintain focus across the imaging area

### Image processing

For all nuclear segmentation, we used custom python scripts to run CellPose (Stringer et al., 2021) in a high throughput manner on DAPI signal. Scripts are available at: https://github.com/arjunrajlaboratory/Trident. Once CellPose masks were obtained, we imported the masks into custom matlab software available at https://github.com/arjunrajlaboratory/dentist2. For human gene RNA FISH, the image analysis pipeline includes thresholding of each fluorescence channel to identify individual RNA FISH spots, connection of RNA FISH spots to a nearest nuclei, and then extraction of spots for all channels and cells. For viral FISH and reporter quantification, we used custom matlab software available at https://github.com/arjunrajlaboratory/dentist2/wiki/Dentist2-Immunofluorescence. Briefly, after importing CellPose masks for segmented nuclei, we drew an annulus around each nuclei to approximate the area of cell cytoplasm. We then calculated the mean fluorescence intensity across the annulus of each nuclei and extracted the data for downstream analysis in R. For automated FISH spot detection of host mRNA, we used custom software available at https://github.com/arjunrajlaboratory/dentist2/wiki/.

### Drug treatment of Calu-3

We used tazarotene (T7080-10MG) at various concentrations to induce *TIG1* expression in Calu-3. For drug treatments, we pretreated cells for 48 hours with either drug or vehicle controls (DMSO) prior to infection. We then collected cells 48 hours post infection and performed single molecule RNA FISH for viral RNA. For each experiment, we also maintained matched mock infected cells (that were not infected) that we subsequently performed FISH on for *TIG1* expression. In order to determine the optimal conditions for *TIG1* induction with tazarotene, we optimized the following parameters; cell passage number post thaw from ATCC (passage 5, passage 6, and passage 7 were tested); the dose of tazarotene (0.3uM, 3uM, 30uM); the time of drug addition (at the time of plating of cells, or one day post cell plating); the surface type used to grow cells (glass wells or plastic tissue-cultureware).

### Bulk RNA sequencing and analysis

We extracted RNA from live cells for bulk RNA-sequencing using the NucleoSpin totalRNA FFPE micro kit (Macherey-Nagel #740969.50) . To prepare sequencing libraries, we used the NEBNext Poly(A) mRNA Magnetic Isolation Module (NEB #E7490L) and NEBNext Ultra II RNA Library Prep Kit for Illumina (NEB #E7770L). To index our samples, we used the NEBNext Multiplex Oligos for Illumina (Dual Index Primers Set 1) oligos (NEB E7600S). All sequencing was performed using an Illumina NextSeq 75 cycle high-output kit (Illumina 20024906) in a paired end format (38 cycles read 1 + 37 cycles read 2). After sequencing, we aligned reads to the human genome (assembly 38; hg38) using kallisto (Bray et al., 2016) and generated count tables with uniquely mapped reads using custom python and R scripts found at https://github.com/arjunrajlaboratory/KallistoSleuth. We then performed differential expression analysis in R v3.6.3 using the limma package and with data from at least 2 biological replicates for each sample and condition.

### Data and Code Availability

All raw and processed data as well as code for analyses in this manuscript can be found at https://www.dropbox.com/sh/1qff4r2v5e7ccdv/AADnfKbB-Wo65ITmS9DBpE99a?dl=0

### Author Contributions

SR, SC, and AR conceived and designed the project with virological and methodological input from SC. SR designed, performed, and analyzed all of the single cell and bulk RNA sequencing experiments including design and generation of all CRISPR-Cas9 cells, supervised by AR. SC designed all virological experiments with JM and SC performing all BSL3 work and JM performing other viral infections. JM performed the qPCR experiments. KA maintained Calu-3 cells for use in all experiments. MCD helped with molecular cloning, single molecule FISH, and image analysis. DS performed the immunofluorescence analysis. NJ helped with single cell RNA sequencing and barcode analysis methods. SR, SC, and AR wrote the manuscript. The authors read and approved the final manuscript.

### Competing Interests

AR receives royalties related to Stellaris RNA FISH probes. All remaining authors declare no competing interests.

## Supporting information

Table S1_CRISPRguides

Table S2_FISHprobes

## Acknowledgements

We thank Sean Whelan (Washington University) for VSV-Spike and members of the Arjun Raj Lab, particularly Yogesh Goyal, Connie Jiang, Benjamin Emert, Ian Dardani, and Phil Burnham for their insightful discussions related to this work. We thank Steph Sansbury of the Ophir Shalem lab for advice on cloning and CRISPR-Cas9 systems. We thank the Genomics Facility at the Wistar Institute, especially Sonali Majumdar and Sandy Widura for assistance with single-cell partitioning and addition of 10x cell identifiers. We thank the members of the Cherry lab, in particular Jorge Acuna and the High-throughput Screening Core for technical and experimental support.

AR acknowledge support from TR01 GM137425, NIH R01 CA238237, NIH R01 CA232256, NIH 4DN U01 DK127405, and NSF EFRI EFMA19-33400. SC acknowledge support from NIH 1-R01-AI-140539, 1R01AI1502461-R01-AI-152362, the Mark Foundation 19-011MIA, and the Burroughs Wellcome Fund. NJ acknowledges support from the Michael Brown Fellowship, NIH T32 GM717044, and NIH F30 HD103378.

