## Supplemental Figures and Captions for "Single cell susceptibility to SARS-CoV-2 infection is driven by variable cell states"

#### Supplemental Figure 1

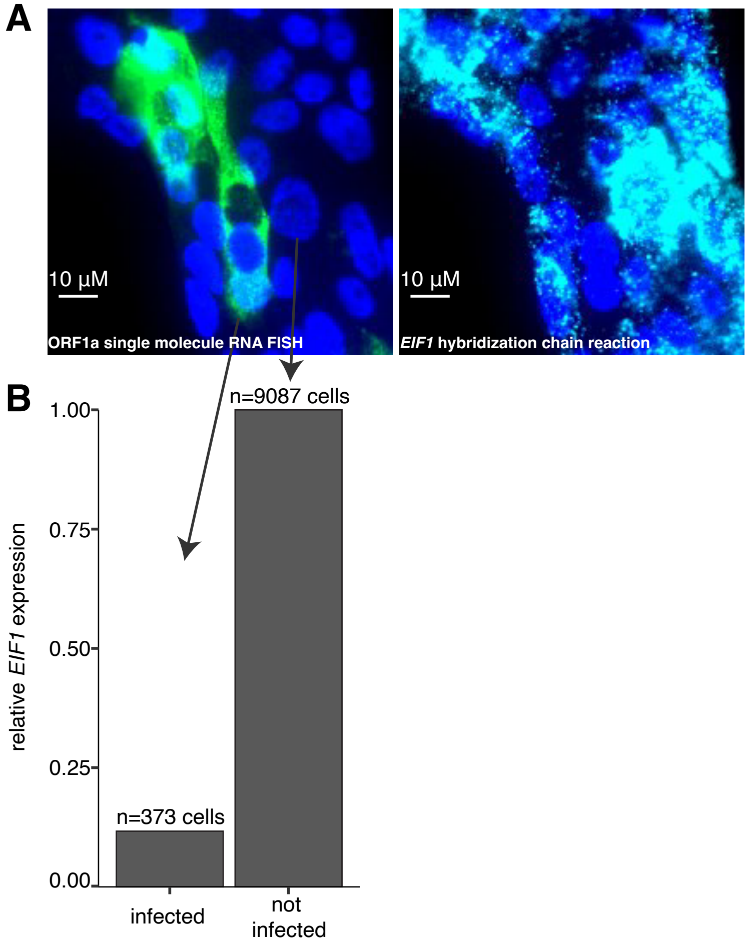

### Supplemental Figure 2

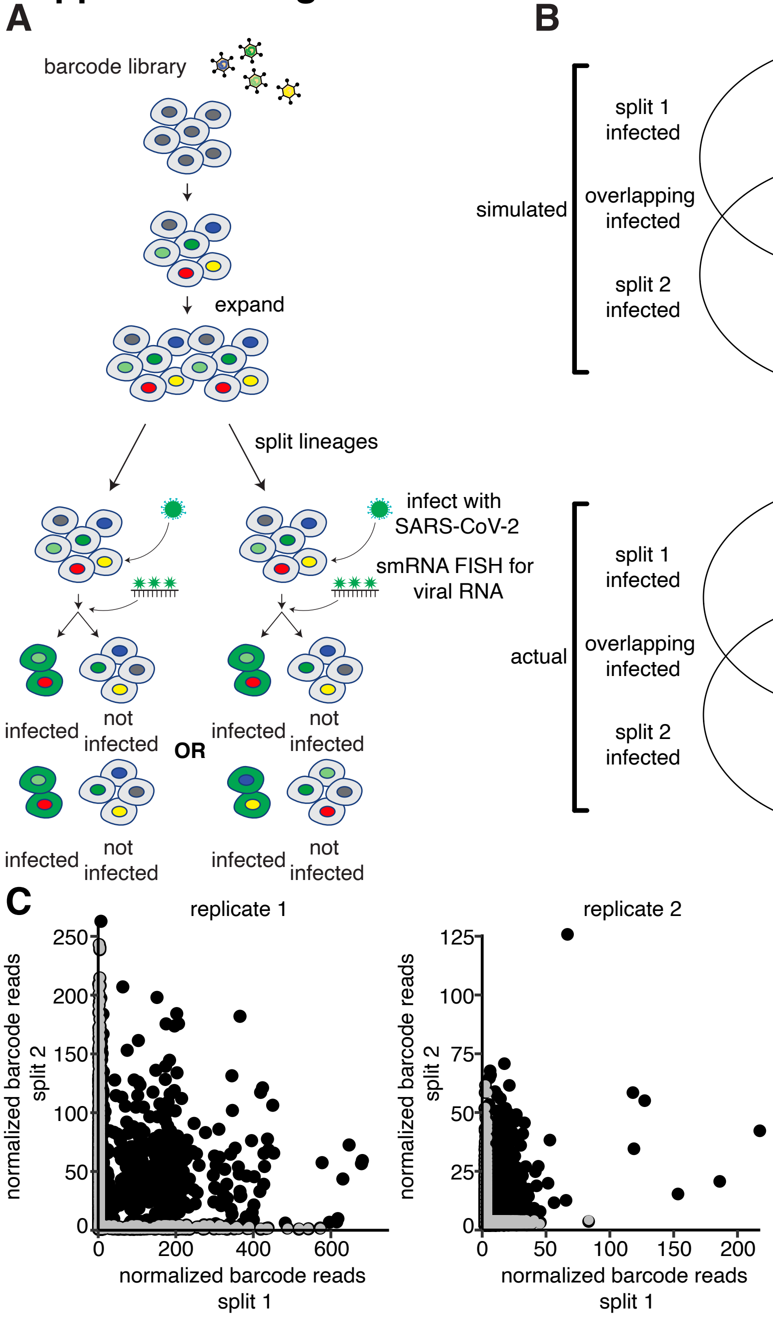

### Supplemental Figure 3

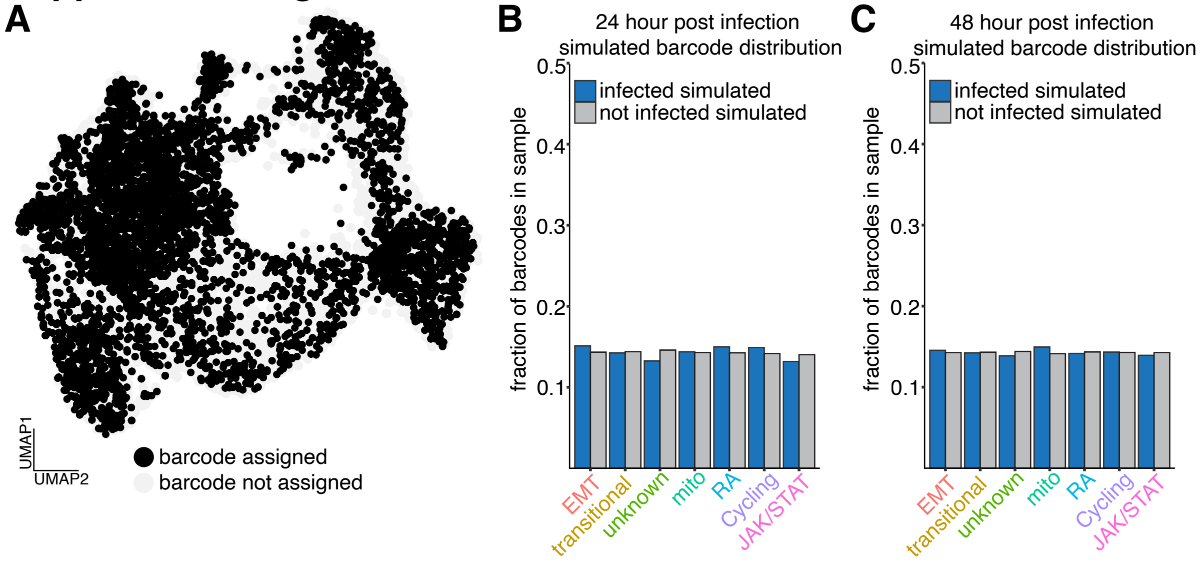

### Supplemental Figure 4

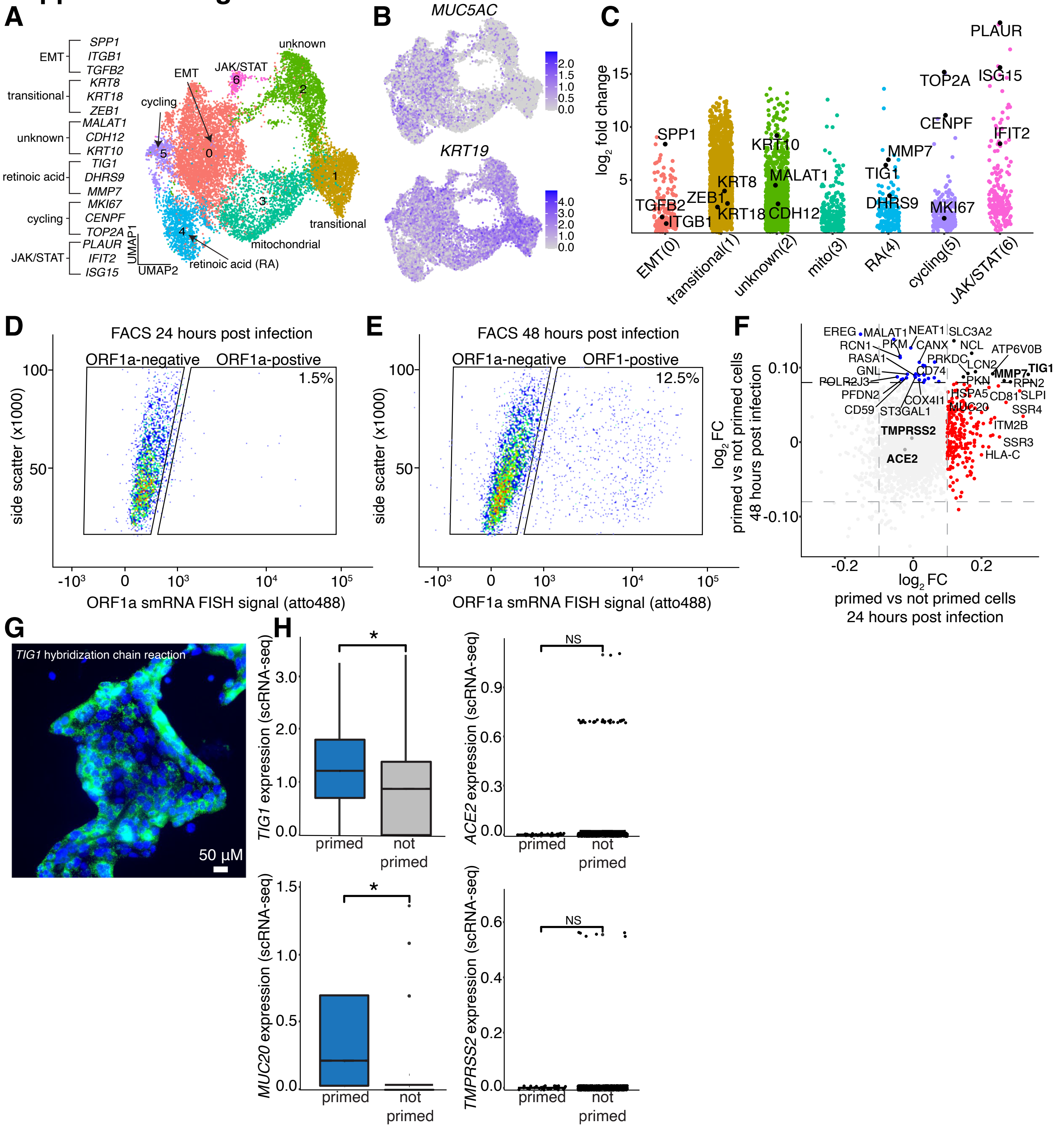

### Supplemental Figure 5

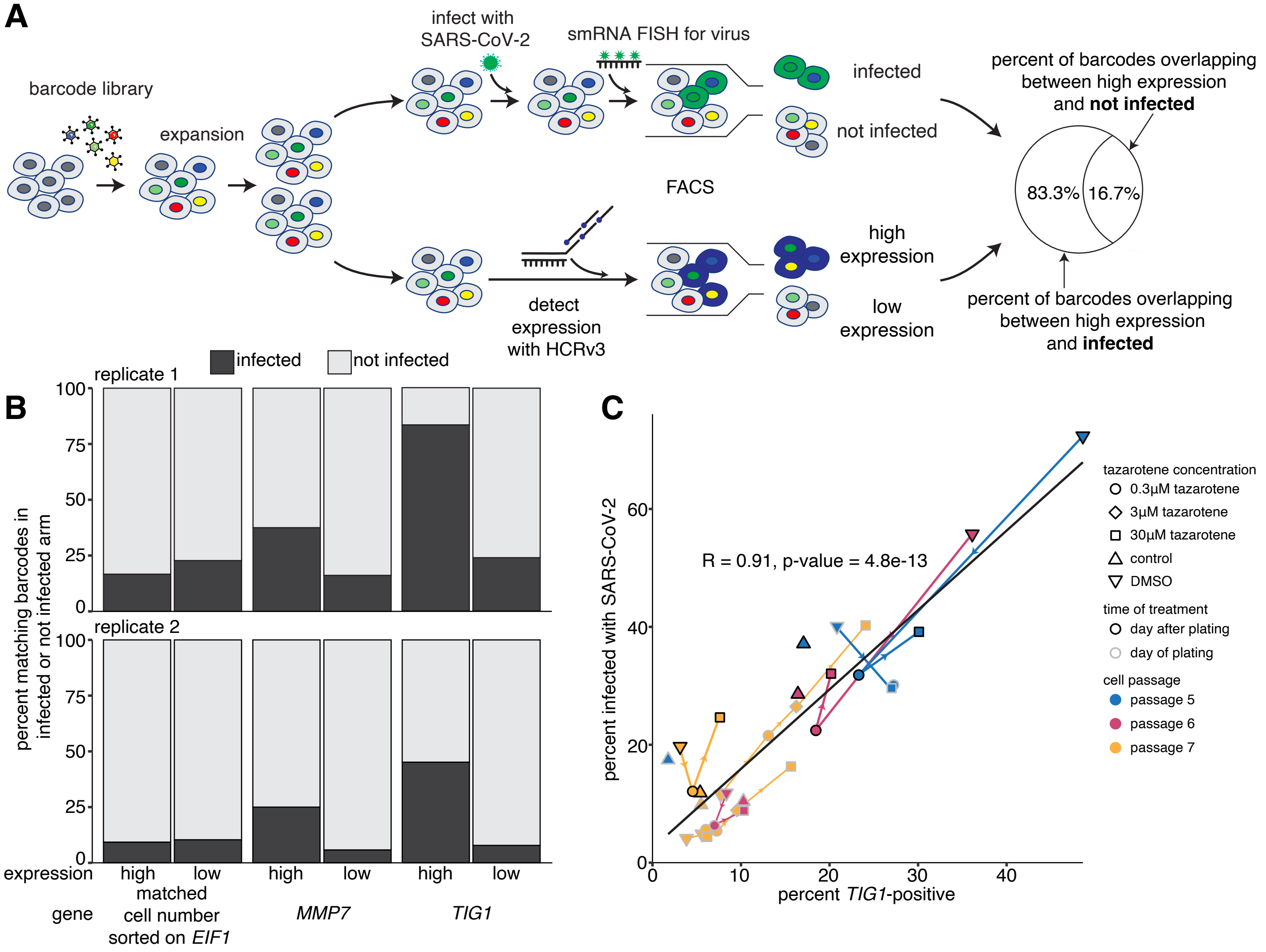

#### Supplemental Figure 6

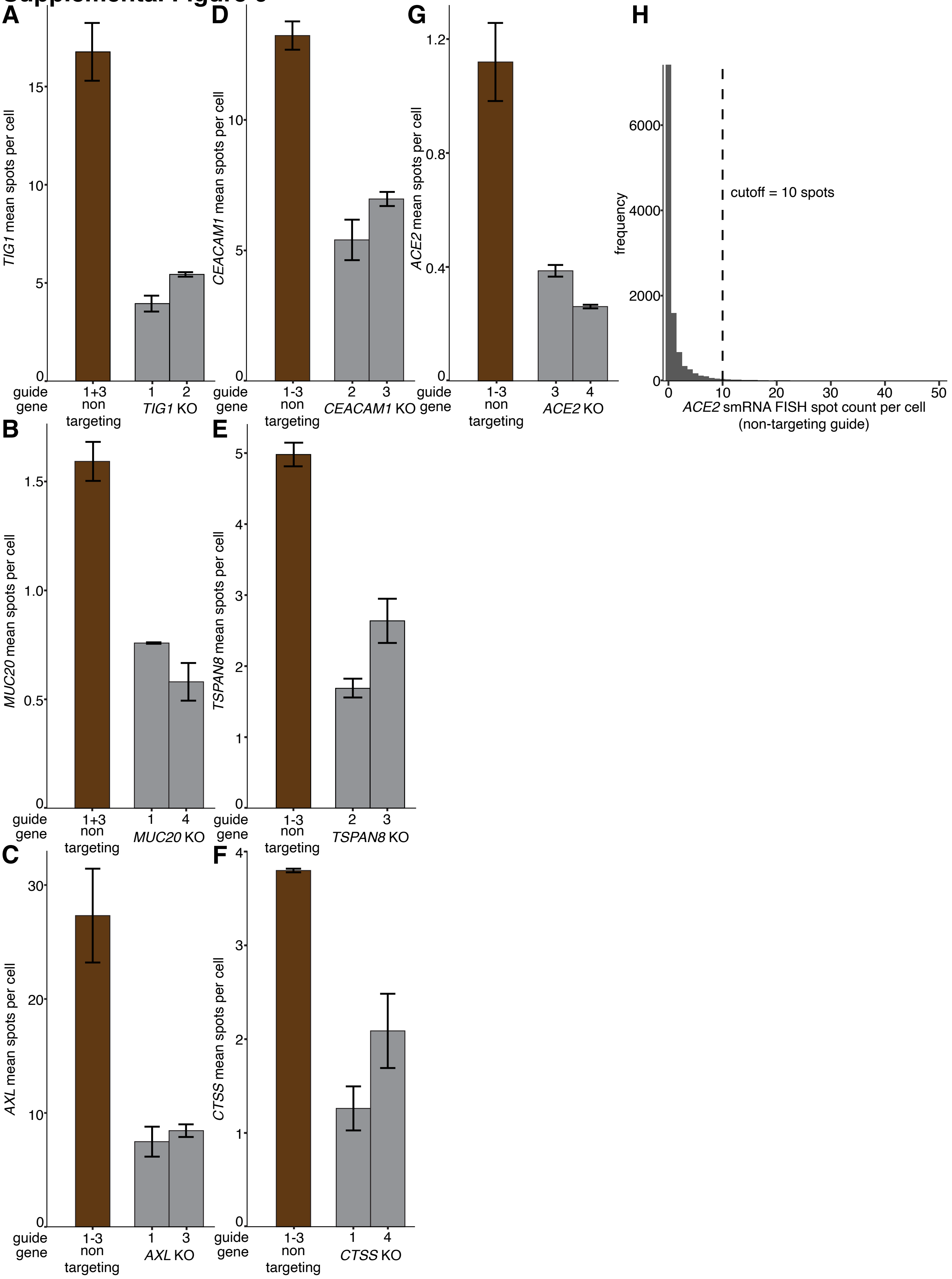

### Supplemental Figure 7

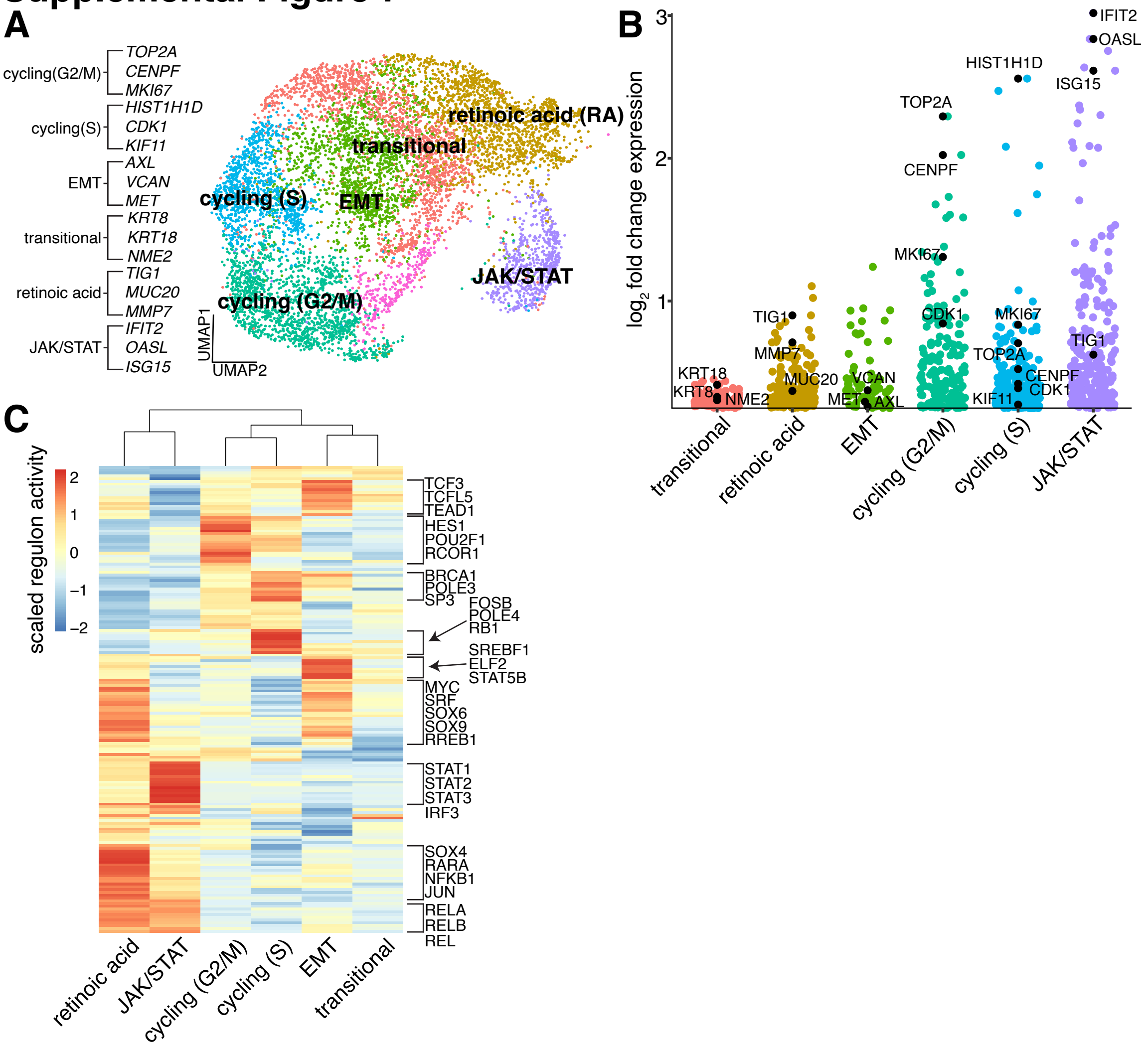

### Supplemental Figure 8

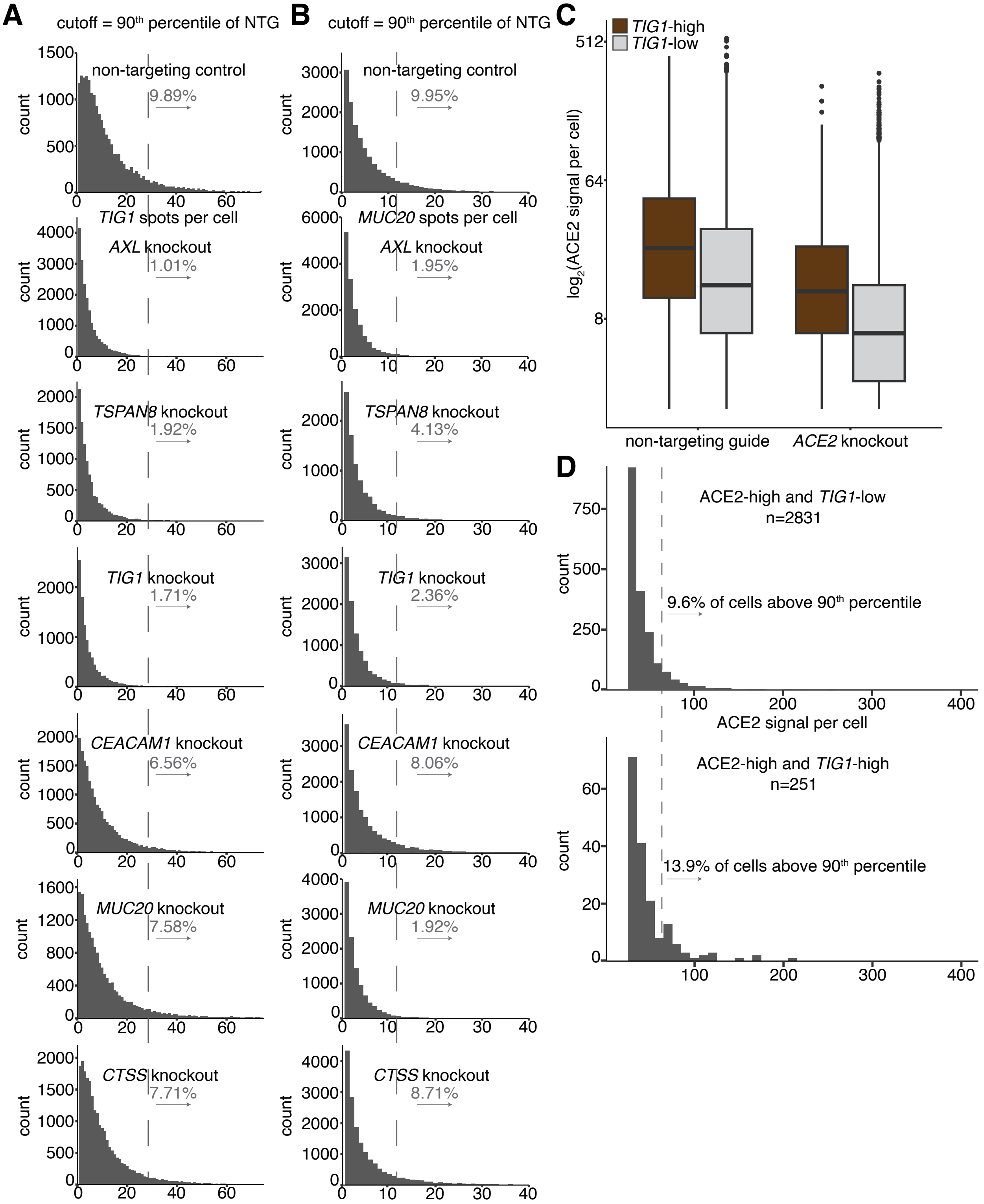

### Supplemental Figure 9

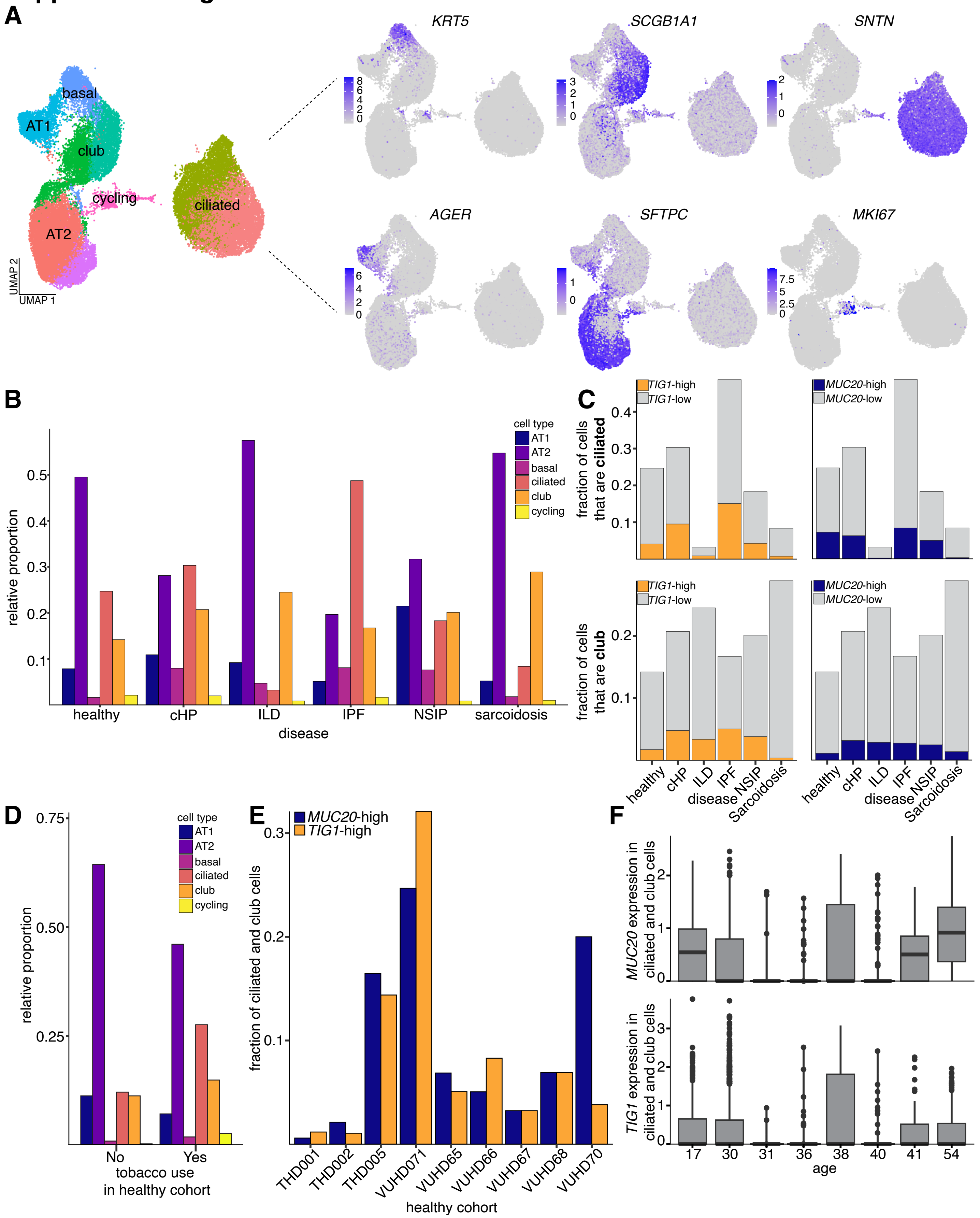

### Supplemental Figure 10

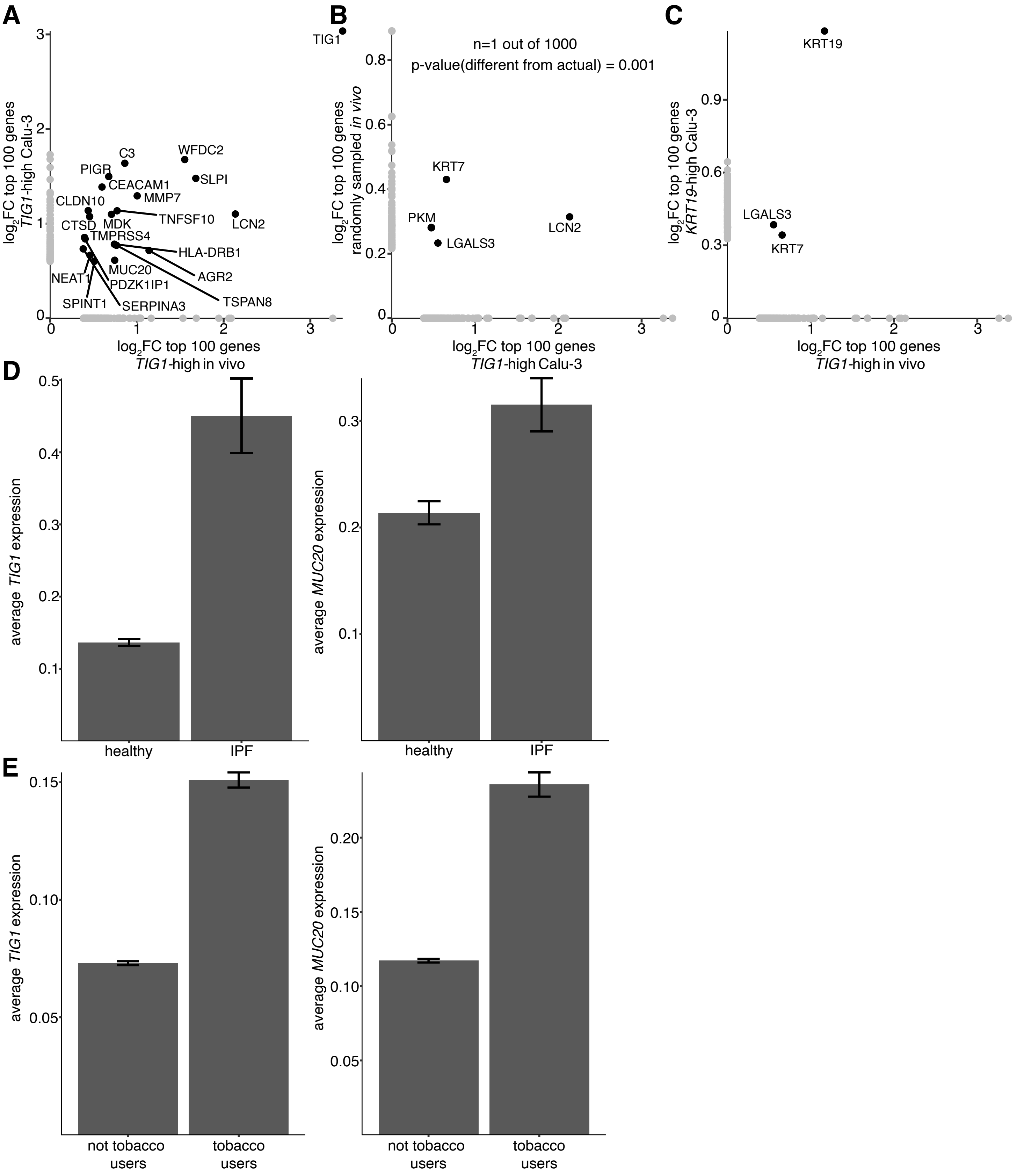

### Supplemental Figure 11

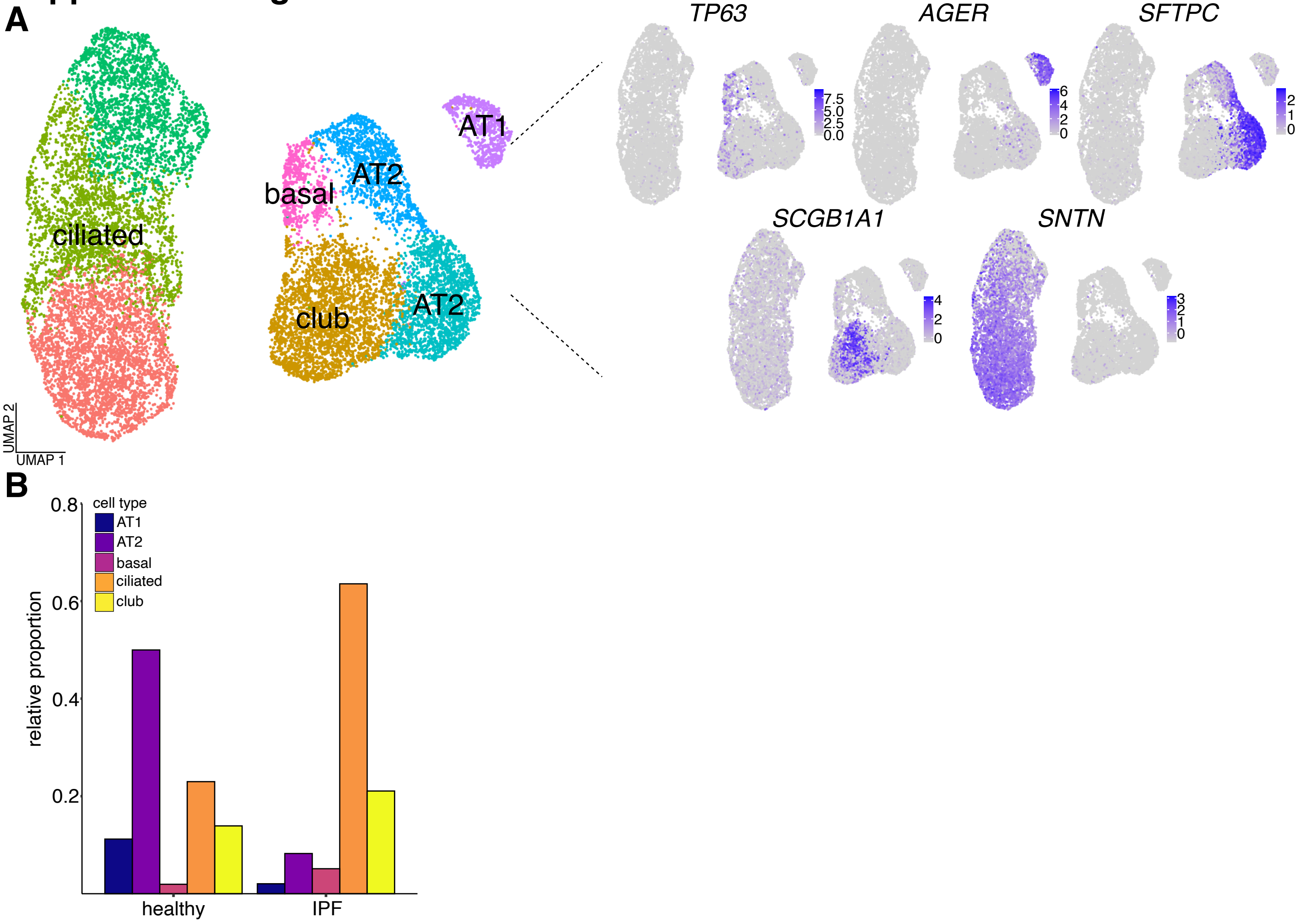

#### Supplemental Figure 12

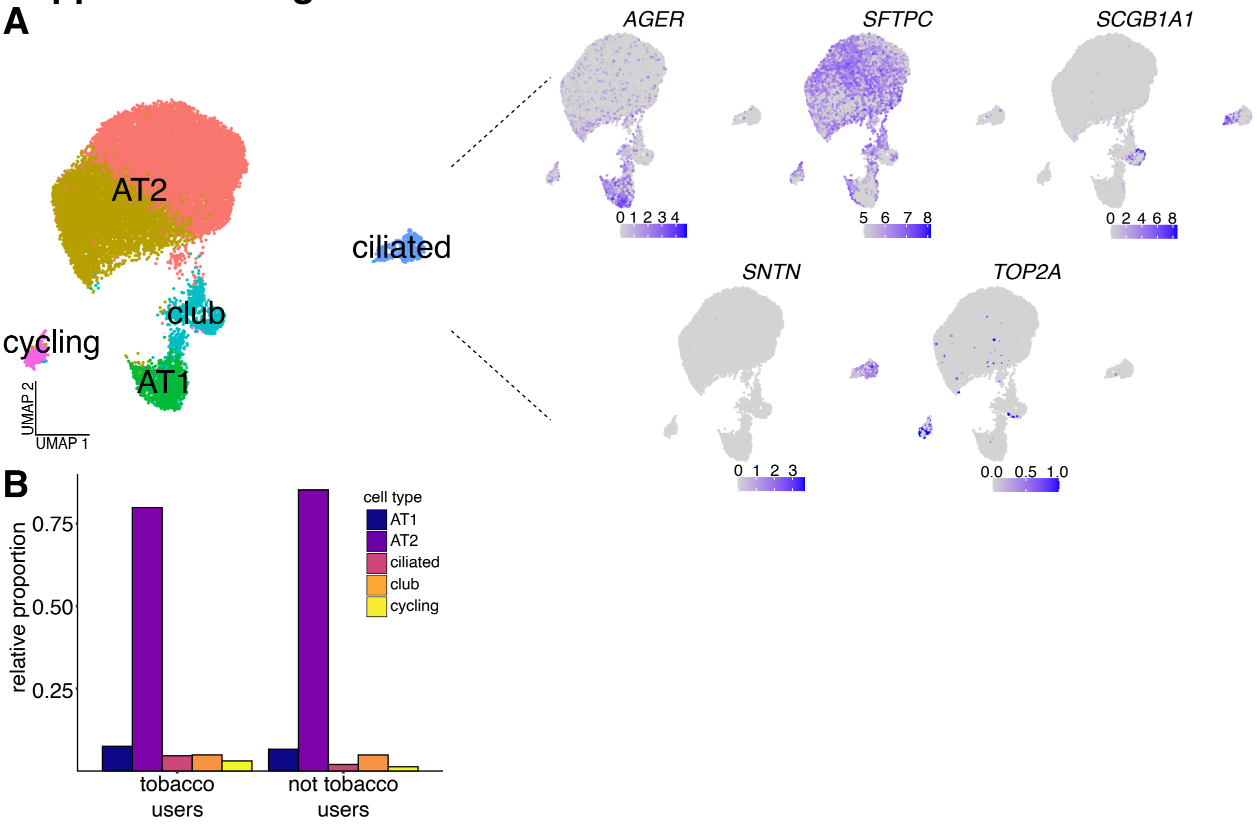

#### Supplemental Figure Captions

**Figure S1:** ***Relative loss of housekeeping gene expression in cells infected with SARS-CoV-2***
**A.** Representative images of cells co-labeled with single molecule RNA FISH against SARS-CoV-2 genome and HCR against the housekeeping gene *EIF1*.
**B.** Relative expression of *EIF1* between infected and not infected cells. Expression was quantified from 373 and 9087 cells from each bin, respectively.

**Figure S2:** ***Quantification of barcode overlap in split clones of Calu-3 demonstrates intrinsic memory for infection***
**A.** Schematic of experimental design to determine if Calu-3 cells have intrinsic susceptibility to SARS-CoV-2 infection. 100,000 Calu-3 were barcoded at a low MOI. After multiple cell divisions (~7 days), clones were split into two arms and infected with SARS-CoV-2. 24 hours post infection, infected cells were fixed, labeled for viral RNA with single molecule RNA FISH, and sorted by FACS. Genomic DNA was extracted to quantify barcode abundance in each split.
**B.** Quantification of barcode overlap across split infections. Top: expected barcode overlap given the number of barcodes introduced and the percentage of cells infected (~2% of total cells) as determined by a barcode recovery simulation (see methods). Bottom: actual barcode overlap between splits, across two biological replicates. In order to assign significance, we performed a one sample z-test comparing the observed barcode overlap in each replicate to the average of the simulated overlap across 1000 repetitions. Four stars refers to a p-value $\leq 0.0001$.
**C.** Correlation of barcode reads across splits for each replicate.

**Figure S3:** ***Recovery and distribution of barcodes in Calu-3***
**A.** UMAP with cells labeled as having a barcode assigned or not. Of n=14,057 cells in the dataset, we were able to reliably assign barcodes to n=5,863 cells (see methods for details).
**B.** To determine the expected number of barcodes recovered in each cluster in UMAP space, we randomly sampled our starting barcode distribution n=100 times into bins that were of the same size as the number of infected and not infected clones recovered 24 hours post infection. After each sampling, we calculated the fraction of barcodes recovered from each cluster, normalized to the size of that cluster. After all sampling was complete, we then calculated the mean and standard deviation for the fraction of barcodes in each cluster for each bin across all simulations.
**C.** We repeated the same sampling procedure as **B**, this time sampling barcodes into bins of the same size as the number of infected and not infected clones recovered 48 hours post infection.

**Figure S4:** ***Rewind identifies distinct transcriptional states that are primed for infection with SARS-CoV-2***
**A.** UMAP of Calu-3 cells subjected to single cell RNA sequencing in the profiling arm of Rewind.
**B.** Normalized expression of *MUC5AC* and *KRT19*.
**C.** Markers of the clusters identified in **A**.
**D.** FACS results after sorting on viral genome single molecule RNA FISH signal in barcoded Calu-3 cells collected 24 hours post infection.
**E.** FACS results after sorting on viral genome single molecule RNA FISH signal in barcoded Calu-3 cells collected 48 hours post infection.
**F.** Differential expression analysis between primed cells and not primed cells from barcodes recovered both 24

hours post infection and 48 hours post infection.
**G.** Expression of *TIG1* as measured by HCRv3 in Calu-3 cells.
**H.** A direct comparison of *MUC20* and *TIG1* normalized expression across primed and not primed cells (significance assigned by performing a wilcox test). We also compared the normalized expression of known host factors required for SARS-CoV-2 infection, *ACE2* and *TMPRSS2*, across primed and not primed cells.

**Figure S5:** ***TIG1 and MMP7 mark a subset of Calu-3 cells that are highly susceptible to SARS-CoV-2 infection***
**A.** We performed an experiment to determine if clones that have high *TIG1* or *MMP7* expression at the time of infection have a greater rate of infection compared to the bulk population. Briefly, we barcoded Calu-3 cells. After a few divisions, we split the barcoded cells into two arms; one of which we infected with SARS-CoV-2. In the other arm, we used HCRv3 to label and then sort clones that had high and low expression of markers (*TIG1*, *MMP7*) identified in primed cells from our Rewind experiment. We defined a cell as high expressing for a given gene if it was in the 90$^{th}$ percentile of signal compared to the rest of the population. All other cells were defined as low expressing. We included a control that was sorted on *EIF1* expression in the same manner as *TIG1* and *MMP7*. In addition, we matched the number of 'high' and 'low' expressing *EIF1* cells recovered to that of *TIG1* and *MMP7*. Of note, the number of high and low *TIG1* and *MMP7* cells were approximately the same. After sorting, we collected genomic DNA from all cells for barcode identification.
**B.** Quantification of barcode overlap between clones expressing high or low amounts of *MMP7* or *TIG1* and clones that are infected. Two independent biological replicates are shown.
**C.** Correlation of the percentage of cells that are expressing *TIG1* to the percentage of cells that are infected with SARS-CoV-2 across a range of drug treatments, passage number, and plating strategies. Lines connect conditions within experiments. Arrows indicate direction of drug concentration increase.

**Figure S6:** ***Validation of gene deletion by CRISPR-Cas9***
**A-G:** single molecule RNA FISH against each target gene for each CRISPR-Cas9 guide tested. Bars represent the average of 2 biological replicates.
**H:** RNA FISH spot count distribution for *ACE2* expression in non-targeting guide cells.

**Figure S7:** ***Gene regulatory networks expressed in Calu-3***
**A.** UMAP representation of additional single cell RNA sequencing data from Calu-3.
**B.** Select markers found to be upregulated in each cluster in UMAP space.
**C.** SCENIC analysis performed on single cell RNA sequencing data to annotate gene regulatory networks present in each cluster in UMAP space**.** Regulon activity has been centered and scaled across clusters for ease of visualization.

**Figure S8:** ***Regulation of the TIG1-high state***
**A:** Distribution of *TIG1* single molecule RNA FISH signal across each knockout tested. We called any cell above the 90$^{th}$ percentile of spot counts positive. For this analysis, we pooled the two guides tested for each gene.
**B:** The same analysis as **A**, but for *MUC20* single molecule RNA FISH.
**C:** Distribution of ACE2 immunofluorescence signal per cell in cells that received either the non-targeting guides or guides against *ACE2*. For this analysis, we pooled the two guides tested for each condition.
**D:** Distribution of ACE2 immunofluorescence signal across cells classified as either ACE2-high and *TIG1*-low or ACE2-high and *TIG1*-high. The cutoff used represents the 90th percentile of ACE2 signal per cell across all

ACE2-high cells, regardless of *TIG1* signal. For this analysis, we pooled the two guides tested for each condition.

**Figure S9:** ***Heterogeneous expression of the* TIG1*-high state is found in vivo***
**A.** Normalized expression of canonical cell type specific markers used to annotate various cell types of the lung epithelium.
**B.** Relative proportion of each cell type identified in **A** across each disease in the dataset.
**C.** Fraction of the total cells in each diagnosis that are ciliated or club. Of that fraction, we calculated the proportion of cells that were *TIG1*-high or *MUC20*-high.
**D.** Relative proportion of each cell type identified in **A** across tobacco users and not users within the healthy cohort of the dataset.
**E.** Fraction of ciliated and club cells that are high for *MUC20* or *TIG1* expression across each patient in the healthy cohort of the dataset.
**F.** Expression of *TIG1* or *MUC20* across age for patients found within the healthy cohort of the dataset.

**Figure S10:** ***Overlap between the* TIG1*-high state identified in Calu-3 cells and the* TIG1*-high state identified within in vivo lung***
**A.** In order to compare the *TIG1*-high state identified in Calu-3 to *in vivo TIG1*-high cells, we looked for overlapping genes within the top 100 differential expressed genes found in each group.
**B.** In order to demonstrate that the overlap found in **A** is representative of a true cell state and not an artifact, we made the same comparison but between randomly sampled cells from *in vivo* lung and *TIG1*-high Calu-3. To assign significance, we repeated this random sampling n=1000 times and then assigned a p-value based on the Permutation test. The data shown are n=1 of the random sampled iterations.
**C.** We also calculated the overlap between differentially expressed genes in Calu-3 cells with high expression of an epithelial cell marker, *KRT19*, to differentially expressed genes found in *TIG1*-high cells *in vivo*.
**D.** Average normalized *TIG1* and *MUC20* expression across patients in the healthy cohort or with IPF. Due to the large variability in the overall number of cells recovered for each patient, a weighted mean and standard error were computed.
**E.** Average normalized *TIG1* and *MUC20* expression across patients who were tobacco users compared to not tobacco users. Due to the large variability in the overall number of cells recovered for each patient, a weighted mean and standard error were computed.

**Figure S11:** ***Breakdown of cell types found across healthy and IPF lung in a second public dataset***
**A.** Normalized expression of canonical cell type specific markers used to annotate various cell types of the lung epithelium.
**B.** Relative proportion of each cell type identified in **A** across healthy and IPF lung.

**Figure S12:** ***Breakdown of cell types found across the lungs of tobacco users and not tobacco users in a second public dataset***
**A.** Normalized expression of canonical cell type specific markers used to annotate various cell types of the lung epithelium.
**B.** Relative proportion of each cell type identified in **A** across tobacco users and not tobacco users.
